# Cerebellar state estimation enables resilient coupling across behavioural domains

**DOI:** 10.1101/2023.04.28.538674

**Authors:** Ensor Rafael Palacios, Paul Chadderton, Karl Friston, Conor Houghton

**Affiliations:** School of Physiology Pharmacology and Neuroscience, University of Bristol, United Kingdom; Wellcome Centre for Human Neuroimaging, Institute of Neurology, University College London, United Kingdom; Bristol Computational Neuroscience Unit, Faculty of Engineering, University of Bristol, United Kingdom

## Abstract

Cerebellar computations are necessary for fine behavioural control and are thought to rely on internal probabilistic models performing state estimation. We propose that the cerebellum infers how states contextualise (i.e., interact with) each other, and coordinates extra-cerebellar neuronal dynamics underpinning a range of behaviours. To support this claim, we describe a cerebellar model for state estimation that includes states interactions, and link the underlying inference with the neuronal architecture and dynamics observed empirically. This is formalised using the free energy principle, which provides a dual perspective on a system in terms of both the dynamics of its physical – in this case neuronal – states, and the inference process they entail. As a proof of principle, we simulate cerebellar-dependent synchronisation of whisking and respiration, which are known to be tightly coupled in rodents. In summary, we suggest that cerebellar-dependent contextualisation of behaviour can explain its ubiquitous involvement in most aspects of behaviour.

## General introduction

Behaviour, defined as the purposeful engagement of a system with its environment, is complex, involving interactions and coordination among different sensory modalities ***Mauk and Ruiz (1992***); ***Fonio et al. (2015***) and executive motor systems ***Bergmann et al. (2022***); ***Warren et al. (2021***); ***Romano et al. (2020***); ***Kurnikova et al. (2017***). The cerebellum is involved in many aspects of behaviour, ranging from stimulus-movement association ***Strata (2015***); ***Fiocchi et al. (2022***) to decision making ***Gao et al. (2018***); ***Deverett et al. (2018***); ***Sendhilnathan et al. (2020***); ***Lin et al. (2020***), and including preparatory activity and movement initiation ***Gao et al. (2018***); ***Li and Mrsic-Flogel (2020***); ***Dacre et al. (2021***); ***Frontera and Léna (2021***), fine tuning of movement trajectories ***Sathyamurthy et al. (2020***); ***Becker and Person (2019***); ***Sauerbrei et al. (2015***), coordination of motor systems and effectors ***Vinueza Veloz et al. (2015***); ***Liu et al. (2020***); ***Romano et al. (2020***), behaviourally relevant perceptual processing ***Baumann et al. (2015***), spatial localisation ***Watson et al. (2019***); ***Rondi-Reig et al. (2022***), and social interactions ***Kelly et al. (2020***); ***Baumann and Mattingley (2022***). Yet, it remains unclear how cerebellar computations underwrite this wide range of functions.

The cerebellum has long been thought to contribute to extra-cerebellar processing by supplying estimates of behaviourally relevant dynamics, based on internal probabilistic (i.e., generative or forward) models ***Paulin (1993***); ***Miall et al. (1993***); ***Wolpert et al. (1998***); ***Liu et al. (2003***); ***Ito (2006***). This view is supported by a large amount of empirical evidence, both in humans ***Therrien and Bastian (2015***); ***Argyropoulos (2016***) and in non-human animals ***Liu et al. (2002***); ***Cerminara et al. (2009***); ***Brooks et al. (2015***), showing that neuronal activity in the cerebellum can read as state estimation ***Tanaka et al. (2019***). However, the question of the exact contribution of cerebellar generative models to extra-cerebellar dynamics remains open.

Three cerebellar features are particularly important for its global involvement in behavioural functions. First, it receives and reciprocates input from most brain regions ***Snider and Stowell (1944***); ***Sobel et al. (1998***); ***Huang et al. (2013***); ***Chabrol et al. (2015***); ***Ishikawa et al. (2015***); ***Wagner and Luo (2020***), making it a locus of convergence for sensorimotor integration. Second, it has a relatively simple network architecture, suggesting that it performs a universal or fundamental computation ***Apps and Hawkes (2009***). Lastly, it is involved in detecting and learning relationships between variables, for example, via cellular plasticity mechanisms ***D’Angelo (2014***), which are essential for optimal behaviour. These features speak to a role of the cerebellum in providing extra-cerebellar regions with estimates of behaviourally relevant states, where, notably, these estimates leverage learned expectations about the interactions and context sensitivity of various latent or hidden states. In other words, assuming that functional behaviour results from the coordination of the dynamics of latent external and somatic states, we propose that the cerebellum is in a key position to efficiently detect, learn and realise context-sensitive interactions among latent states, in order to constrain and orchestrate extra-cerebellar dynamics.

This description of cerebellar computations can be formalised within computational neuroscience in many ways. For example, as a Smith predictor ***Miall et al. (1993***), as a forward model, as model predictive control, as an instance of predictive coding and so on ***Doya (1999***); ***Frens and Donchin (2009***); ***Friston and Herreros (2016***); ***Koziol et al. (2014***); ***Paulin (2005***); ***Ramnani (2014***); ***Shadmehr and Krakauer (2008***); ***Tseng et al. (2007***). We will adopt a generic formalism provided by the free energy principle (FEP) that encompasses all of these accounts. The FEP furnishes an account of selforganisation based on minimising prediction errors (a.k.a, surprisal), or, equivalently, maximising model evidence (a.k.a. marginal likelihood) ***Friston (2009***, 2013). This formulation of self organisation as an optimisation process describes how such a system persists over time by maintaining a separation between itself and the environment, while interacting with that environment.

Crucially, the FEP offers a dual perspective on the dynamics of a system ***Ramstead et al. (2022***), linking its trajectory in the space of physical – in this case neuronal – states to a trajectory in an implicit encoding space, where each point parameterises a probability distribution over external or latent states. As a result, a system successfully interacting with its environment is necessarily a good probabilistic model of this environment (see the narrative summary in Methods and Materials for more details). This physics of self-organising systems can be applied to neuronal networks like the cerebellum ***Palacios et al. (2019***); ***Isomura (2022***), therefore providing a principled way to describe how the cerebellum implements probabilistic models to support behaviour: in essence, under the FEP, the cerebellar architecture defines the form of an implicit generative model, and neuronal dynamics reflect an inference process based on this model. Technically, neuronal dynamics become a gradient flow on variational free energy, where (negative) variational free energy scores the evidence for a generative model of the world at hand.

The application of the FEP to functional brain architectures generally reduces to identifying the structure and functional form of the generative model that best explains neuroanatomy and neurophysiology. The hypothesis in this work is that the cerebellum occupies a *deep* or alternatively high level in a hierarchical generative model of sensorimotor exchange. This might seem slightly counterintuitive, given the intimate connectivity of the cerebellum with the brainstem; however, the empirical properties of the cerebellum described above speak to a high-level role in contextualising lower, extracerebellar, levels. This can be abduced from the widespread connectivity of the cerebellum to other regions and from lesion-deficit models: for example, cerebellar lesions lead to deficits such as cerebellar ataxia that are characterised by a loss of coordination as opposed to movement *per se*. Structurally, this view means that the cerebellum generates the context for, but not the content of, sensorimotor coordination.

From the perspective of active inference, this means the cerebellum has to recognise the ongoing context or mode of behaviour – through assimilating ascending prediction errors from extra-cerebellar levels – while, at the same time, supplying empirical priors to guide or constrain (i.e., contextualise) dynamics at lower levels via descending predictions to extra-cerebellar levels. Here the context is determined by control states or parameters, whose inference depends on the co-occurrence or joint evolution of discrete behavioural states – the content. These control states in turn enter in a nonlinear fashion to modulate the dynamics of lower levels. This requires a highly nonlinear extra-cerebellar model, in contrast to a simpler, weakly nonlinear model in the cerebellum, reflected in its simple circuitry. In what follows, we will use a linear approximation to cerebellar dynamics and nonlinear (Kuramoto) oscillators at the extra-cerebellar level that are coupled via a cerebellar control state. This kind of architecture undergirds many active inference simulations of coordinated behaviour and speaks to a separation of context and content in deep generative models of coordination dynamics, see for example, ***Friston and Kiebel (2009***); ***Jirsa and Kelso (2005***); ***Kelso (2021***); ***Kiebel et al. (2008***, 2009).

This paper is organised as follows: we first introduce the FEP, as the theoretical framework for active inference and learning in the brain. Next, we describe how cerebellar models could contribute to active inference by outlining the functional form they could take, and establish a mapping from elements of the model to components of the cerebellar circuitry. We then simulate inference in a cerebellar model during motor coordination of respiratory and whisker behaviour. The results of the simulations reveal that cerebellum provides a corrective coupling between the respiratory and whisking networks to maintain synchronisation in the presence of different types of noise and environmental perturbation. These numerical studies are intended to showcase how cerebellar-based state estimation might contribute to active inference, and ensuing action, in extra-cerebellar structures. Finally, we discuss the implications of cerebellar models for simulations of whole-brain dynamics.

### Neuronal dynamics as inference

Under the FEP, neuronal dynamics are read as an inference process, which results in goal-directed behaviour ***Friston et al. (2016***); ***Da Costa et al. (2021***). In this picture, environmental states *ϑ* cause observations *y*, and inference serves to estimate *ϑ*. Inference is based on a generative model *m*

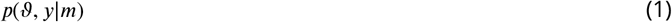

of how *ϑ* relates to *y*; the form of this model is defined by the architecture of the underlying neuronal network. Mathematically, neuronal inference implies that beliefs about environmental states, encoded by the activity of neuronal populations, will evolve to match the posterior probability over external states:

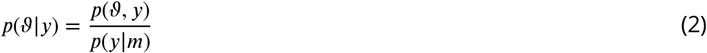

Since the denominator in this equation requires integration over all possible states, it is often intractable. Thus, the FEP proposes that the brain can be viewed as performing approximate Bayesian inference, where the negative logarithm of *p*(*y*) is approximated by an upper bound called variational free energy (*F*). By assuming that the brain encodes approximate beliefs, *q*(*ϑ*), about external states and in particular their most likely or expected value *μ*_*ϑ*_, the variational free energy *F* scores the (negative log) evidence ***Buckley et al. (2017***):

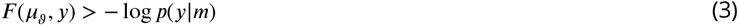

This quantity is a function of the model entailed by the network, while expectations about hidden states are encoded by its activity, and available observations ***Friston (2009***, 2010). Crucially, the expectation *q*(*ϑ*) that minimises *F*, or maximises model evidence *p*(*y*|*m*), is also the one that best approximates the expectation of the posterior *p*(*ϑ*|*y*) (see Eq. 14 in methods).

Intuitively, given a certain model *m, F* is minimised by maximising the probability of co-occurrence of expected states *μ*_*ϑ*_ and observations *y*. This, in turn, can be achieved either by changing *μ*_*ϑ*_, making *q*(*ϑ*) a better approximation of *p*(*ϑ* |*y*), or by changing *y* by acting on external or latent states so that they conform to internal expectations, hence increasing model evidence *p*(*y*|*m*). This process is at the core of active inference, an application of the FEP that formalises the action-perception cycle in terms of Bayesian inference. Note that under simplifying assumptions about random fluctuations the free energy gradients that drive internal dynamics and action can be expressed as prediction errors. This leads to a description of neuronal dynamics in terms of predictive coding or Bayesian filtering, while action can be reduced to simple reflex arcs that minimise proprioceptive prediction errors, in the spirit of the equilibrium point hypothesis or, on another view, perceptual control theory, see for example ***Feldman and Levin (1995***); ***Friston et al. (2011***); ***Mansell (2011***).

### Cerebellar contribution to neuronal inference

The ability of a system to minimise *F* depends on how well its implicit generative model is able to track real environmental dynamics, or alternatively make it match its own dynamics, by maximising model evidence – sometimes known as self evidencing ***Hohwy (2016***); ***Palacios et al. (2020***). When considering cerebellar computations, one hypothesis is that they improve state estimation or inference in other brain regions to afford more efficient inference. This could be achieved by rapidly contextualising extra-cerebellar inference; that is, the cerebellum might provide estimates about what should be inferred by extra-cerebellar structures given a specific context, where the latter corresponds to what is simultaneously being estimated in other brain regions.

Contextualisation is needed because when a system interacts with its environment, behaviour requires coordination; consequently, neuronal dynamics encoding and controlling external and somatic states – across brain regions – must be coordinated. Clearly, extra-cerebellar regions can coordinate by interacting with other regions through reciprocal connections. However, this stream of information, although necessary, might be relatively slow and computationally expensive, due to intrinsic (axonal) delays in communication and because of the complexity and diversity of the underlying models and ensuing neuronal inference processes.

The cerebellum, in contrast, is able to simultaneously integrate information from different brain regions, and its implicit generative model is general: simple, yet powerful enough to efficiently estimate states that are conditioned on each other. In this view, the cerebellum contextualises behaviour by efficiently learning and realising the necessary coordination among states, by predicting, and therefore generating, smooth, synchronous and concerted dynamics. This is illustrated in Fig. 1, where the cerebellar model is placed on top of a modular generative model implemented by extracerebellar structures. Crucially, the advantage of this contextual inference relies on how well the cerebellum can bind together brain dynamics, which in turn depends on the underlying generative model.

**Figure 1.**
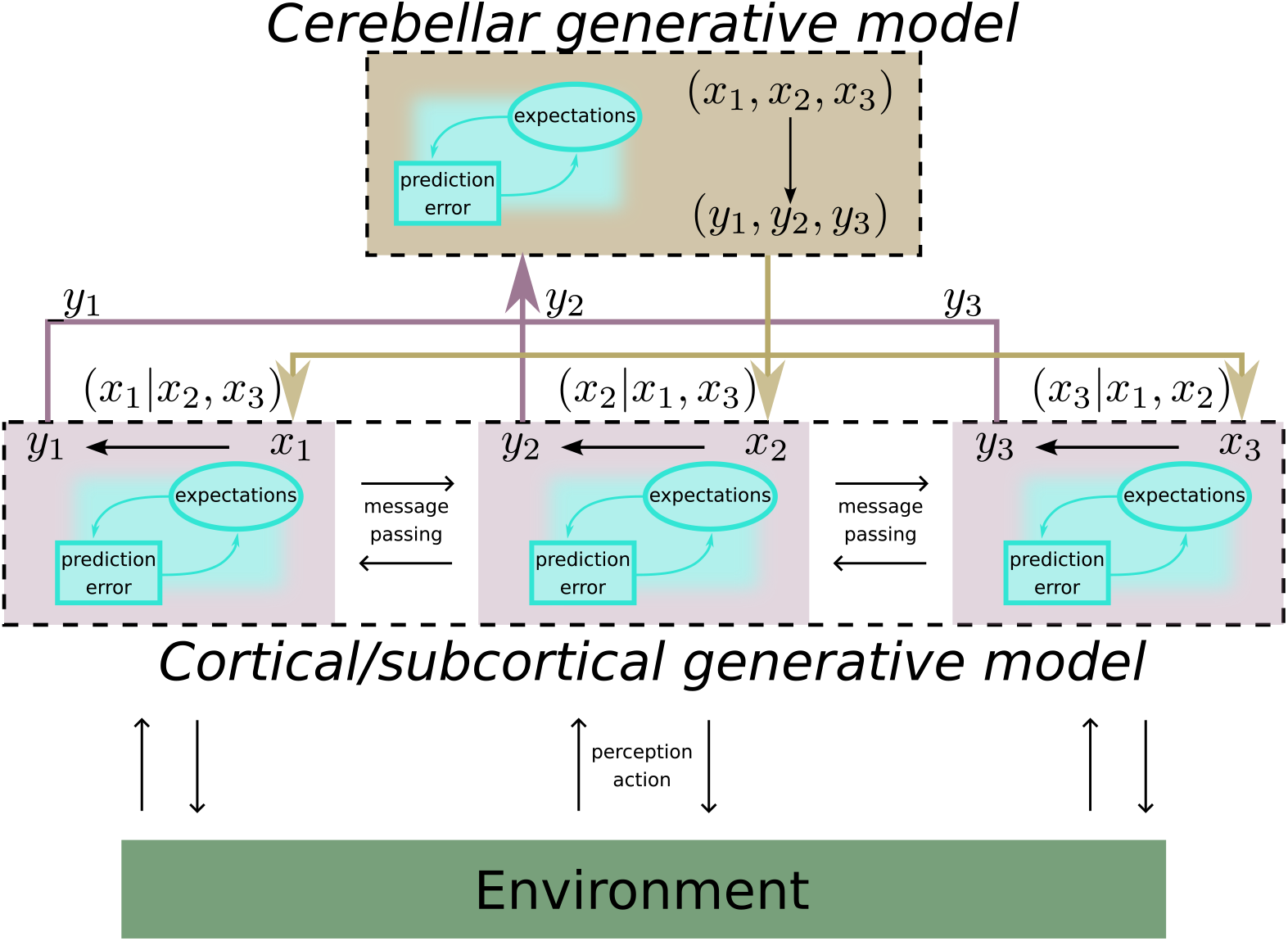
An illustration of cerebellar contribution to active inference in the brain. Extra-cerebellar structures implement a complex generative model of the environment, exemplified by three modules representing different streams of information (violet boxes), all potentially involving hierarchical processing. These modules can underpin discrete but interlinked behaviours, such as whisking and respiration. Within each module, neuronal populations infer hidden state *x* from observation *y* based on forward models (black arrows) of how *x* generates *y*. The accompanying inference process drives neuronal dynamics to update expectations about *x* by minimising prediction errors. Furthermore, modules can communicate with each others via within-module message passing, involving predictions and prediction errors. The cerebellum, on the other hand, concomitantly receives and integrates information (observations or prediction errors) from many brain regions, and implements a forward model of how *x*′s interact at lower (extra-cerebellar) levels (black line from all *x*′s to all *y*′s), which in turn will inform cerebellar estimates of latent states or modes of coordination. Consequently, as the cerebellum both recognises and predicts extra-cerebellar dynamics, it can realise expected interactions by providing top-down (empirical prior) constraints on extra-cerebellar inference.

### The cerebellum: internal model and neuronal dynamics

We suggest that the relatively homogeneous and simple cerebellar architecture implements a state space model. This model includes observations ***y***, carrying information from somatic or extra-cerebellar structures to the cerebellum, hidden states ***x***, corresponding to behaviourally relevant states that need to be inferred, and hidden causes ***v***, which are latent control states mediating interactions among hidden states. (Bold face refers to vectors.) This cerebellar model assumes that ***y*** are generated from ***x***, which in turn have dynamics controlled by ***v*:**

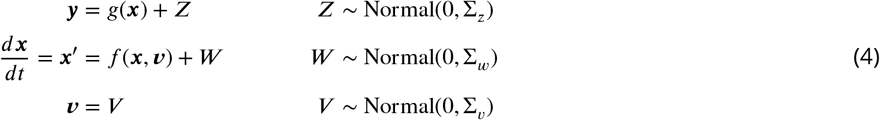

A key feature of this model is that it embodies assumptions about how ***x*** changes over time, allowing the cerebellum to extract information about the trajectory (temporal derivatives) of hidden states ***x*** from streams of observations ***y***; please see below for a discussion of how this could be implemented biologically. Moreover, the mapping from latent states to observations and equations of motion can be approximated with a linear form, emulating the simplicity of the cerebellar circuitry:

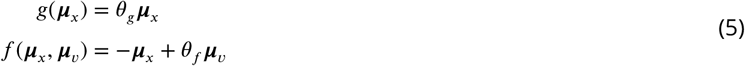

These two equations, Eq. 4 and Eq. 5, describe an *implicit* cerebellar model of how ***x*** and ***v*** generate ***y***, in the sense that the computations carried out by the underlying neuronal dynamics underwrite its inversion: the inference process mapping from consequences, ***y***, to causes, ***x*** and ***v***. In more detail, with this particular model in place, state estimation reduces to finding the expected values ***μ***_***x***_, expected motion ***μ***_*x*′_, and ***μ***_*v*_ that best explain observations ***y***. Given the generative model above, we can specify the requisite estimation equations for ***μ***_*x*_, ***μ***_*x*′_ and ***μ***_*v*_ as follows:

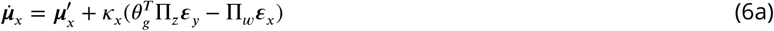

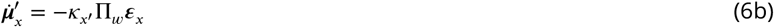

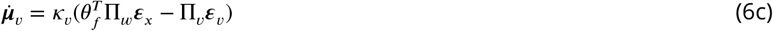

where the *k*′s are learning rates and ***ε***′s are prediction errors:

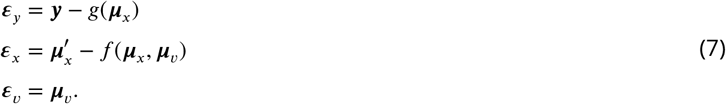

We can now map the inversion dynamics described in Eq. 6 and Eq. 7 onto neuronal components and dynamics of the cerebellum. Specifically, we associate observations *y* with mossy fibre (**mf**) input, hidden causes ***μ***_*v*_ with Purkinje cell population (**Pjc**) activity, and assume that both hidden states ***μ***_*x*_ and velocities ***μ***_*x*′_ are represented by the granule cell population (**grc**) (see Figure 2). Therefore, **grc** activity reflects not only the instantaneous value of hidden states ***μ***_***x***_, but also their change ***μ***_*x*′_, and potentially higher order derivatives, effectively encoding the trajectory of latent states as the coefficients of a Taylor expansion of the path as a function of time.

**Figure 2.**
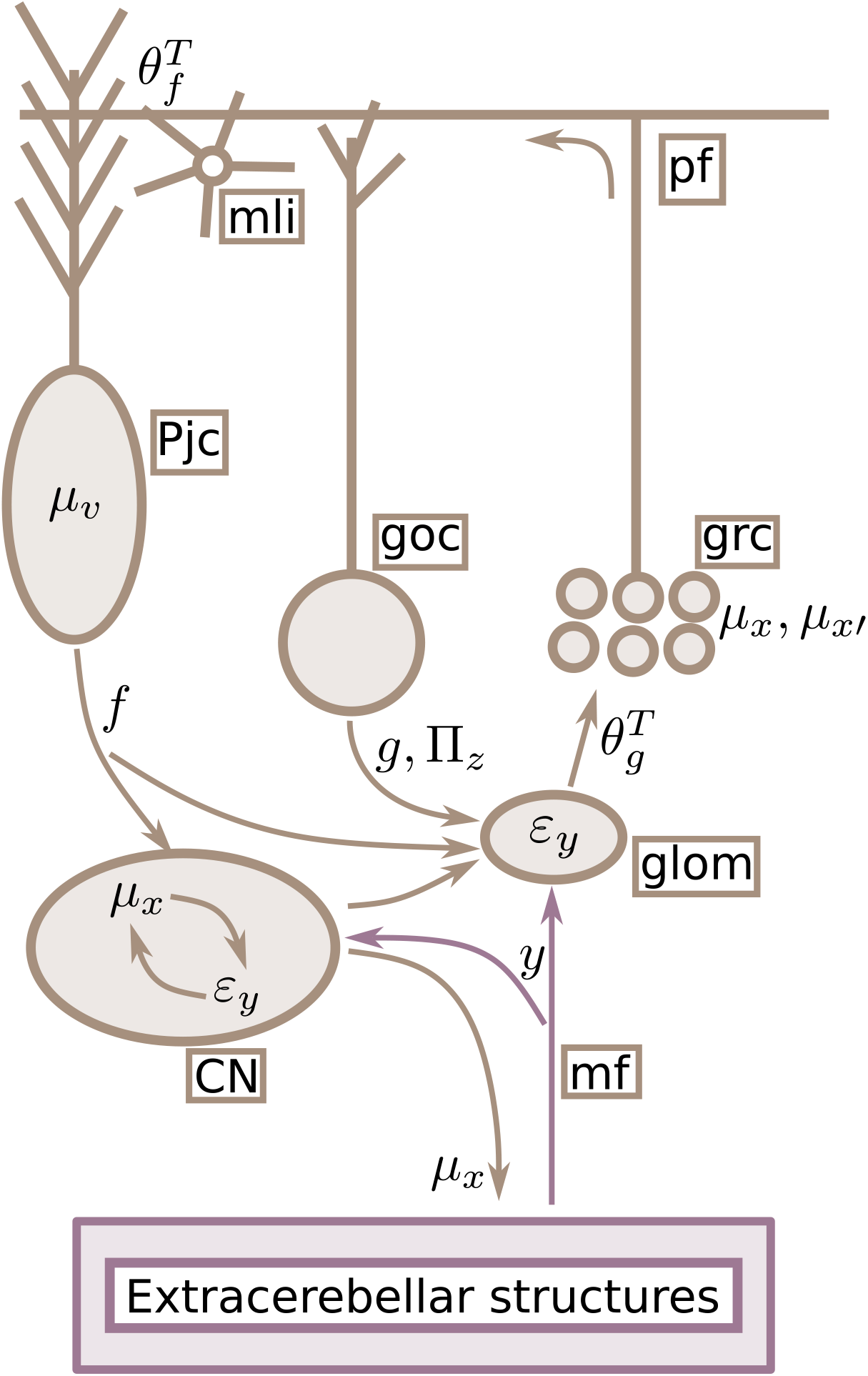
Mapping elements of the model to cerebellar circuitry. Neuronal and model elements are highlighted in brown and black, respectively. The cerebellum receives observation *y* from extra-cerebellar structures through **mf**, which drive prediction errors ***ε***_*y*_. This information enters the **CN** as well as the **glom** – specialised structures in the cerebellar cortex where the excitation-inhibition balance (E-I) drives **grc** and **Goc** activity. E-I results from the integration of different inputs, including **Goc, CN** and **Pjc** activity. Computationally, this is where feedback predictions (*g*, from **Goc**, *f*, from **Pjc** and **CN**) minimise ***ε***_*y*_ and precision (Π_*z*_, from **Goc**) modulates it. The residual ***ε***_*y*_ then drives **Pjc** and **mli** activity via **pf**, based on the particular connectivity encoded by 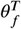 ; this connectivity encodes learned interactions among latent states, which are thus reflected in the dynamics of control states *μ*_*v*_ (**Pjc** activity). These hidden causes are then the source of predictions *f* modulating **CN** and (directly and indirectly) **grc** activity. This modulation constrains estimates of latent states *μ*_*x*_ according to expectations about their interactions. Finally, these estimates couple back to extra-cerebellar models via **CN** collaterals, updating estimates therein. Mossy fibres **mf** ; Purkinje cells **Pjc**; granule cells **grc**; Golgi cells **goc**; glomeruli **glom**; cerebellar nuclei **CN**; molecular layer interneurons **mli**.

This temporal property is made explicit in the equation for 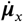, Eq. (6a), where changes in expected values of hidden states, 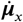, follow ***μ***_*x*′_. From a biophysical standpoint, this means that the recent neuronal history impacts current state estimation ***Hansel et al. (2001***); ***Straub et al. (2020***). Additionally, 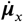 also depends on 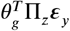 and Π_*w*_*ε*_*x*_: the former equates to new or unexplained information ***ε***_***y***_, based on observations *y* and predictions *g*(***μ***_***x***_), where ***ε***_***y***_ can be thought of as the net input to **grc** computed within glomeruli (**glom**); a specialised structure containing dendritic and axonal terminals from **grc**, Golgi cells (**goc**) and **mf**.

This net input drives state estimation depending on the connectivity matrix 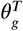 involving **glom** and **grc**, and is weighted or contextualised by its precision 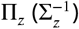 via **goc** inhibition, setting **grc** response gain and threshold ***Palacios et al. (2021***); this weighting by Π_z_ controls how much **mf** drives the network, based on context-dependent precision of incoming information. On the other hand, the last term, Π_*w*_***ε***_*x*_, incorporates feedback from **Pjc** representing hidden causes ***μ***_***v***_. This contribution may occur either via recurrent inputs from cerebellar nuclei (**Cn**) ***Houck and Person (2015***); ***Ankri et al. (2015***) – whose dynamics are driven by **mf** and shaped by **Pjc** inhibition – or through direct **Pjc** modulation of cerebellar cortical interneurons ***Witter et al. (2016***) and grc ***Guo et al. (2016***).

Next, the 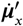 equation, Eq. 6b, specifies updates of encoded velocities, 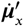: these updates depend on predictions *f* via ***ε***_*x*_, and in absence of (**mf**) input ultimately attract ***μ***_***x***_ (**grc** activity) toward zero (enforcing sparsity). Similarly to 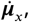, updates 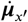 are a function of ***μ***_***v***_, and could plausibly rely on the same feedback mechanisms involving **Pjc** and **Cn**. Interestingly, one could also endow the generative model in Eq. (4) with the ability to ‘observe’ changes in ***y***, namely, 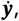, in addition to their instantaneous value. Although, here, we consider the simplest generative model, this addition would effectively allow the history of **mf** input to play a direct role in **grc** dynamics; for example via short-term plastic changes in **mf** boutons ***Xu-Friedman and Regehr (2003***). The ensuing prediction error would then enter the updates of ***μ***_*x*′_, increasing the accuracy of estimates.

Lastly, the equation for 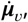, Eq. (6c), describes ***μ***_***v***_ (**Pjc** firing) updates that rely on ***μ***_***x***_ and ***μ***_*x*′_ (**grc** activity) via ***ε***_*x*_; crucially, how this information is transmitted depends on 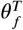, which we associate with the particular connectivity afforded by parallel fibres (**pf**), terminating on molecular layer interneurons **mli** and **Pjc**.

Finally, the cerebellar model is coupled to extra-cerebellar regions because it sends back estimates of latent states, ***μ***_***x***_; these are constrained by cerebellar expectations about their interactions or associations, encoded in the **pf** connectivity structure, 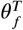. This coupling is mediated by **Cn**, whose activity also encodes hidden states ***μ***_***x***_, and are assumed to receive the same information as cerebellar cortex via **mf** collaterals.

### Simulation of motor coordination

Here, we substantiate our formulation of cerebellar computations with simulations of motor coordination. We start from the basic idea that integration of information across multiple brain regions is crucial for coordinated behaviour, and that the cerebellum contributes to this functional integration by learning to infer interactions between sensory modalities, effectors and task contingencies (e.g. preferred outcomes or reward). A key mechanism through which interactions are learned is synaptic plasticity at the level of **pf** terminals, as in the case of learning temporal associations between conditioned and unconditioned stimuli ***Coesmans et al. (2004***) or motor adaptation during eye movements ***Yang and Lisberger (2013***). Here, we show how acquired expectations about motor coordination, encoded in the **pf** connectivity matrix 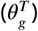, can guide dynamics in extra-cerebellar structures, as the cerebellum provides them with top-down (prior) constraints, hence ensuring optimal behaviour.

In particular, we focus on the coordination of the respiratory and whisker system in mice, which underwrites exploration of the environment in rodents ***Kurnikova et al. (2017***) (Figure 3). The production of these behaviours is underpinned by a rich network of interconnected brain structures, including brainstem pattern generators as well as premotor cortical areas, many of which also receive projections from **CN *Arshavsky et al. (1997***); ***Bellingham (1998***); ***Romano et al. (2020***); ***Takatoh et al. (2022***); ***Golomb et al. (2022***). This richness hints at the complexity of the extra-cerebellar model controlling orofacial behaviour in rodents, to which a key cerebellar contribution appears to be the coordination of its various components ***Romano et al. (2020***).

**Figure 3.**
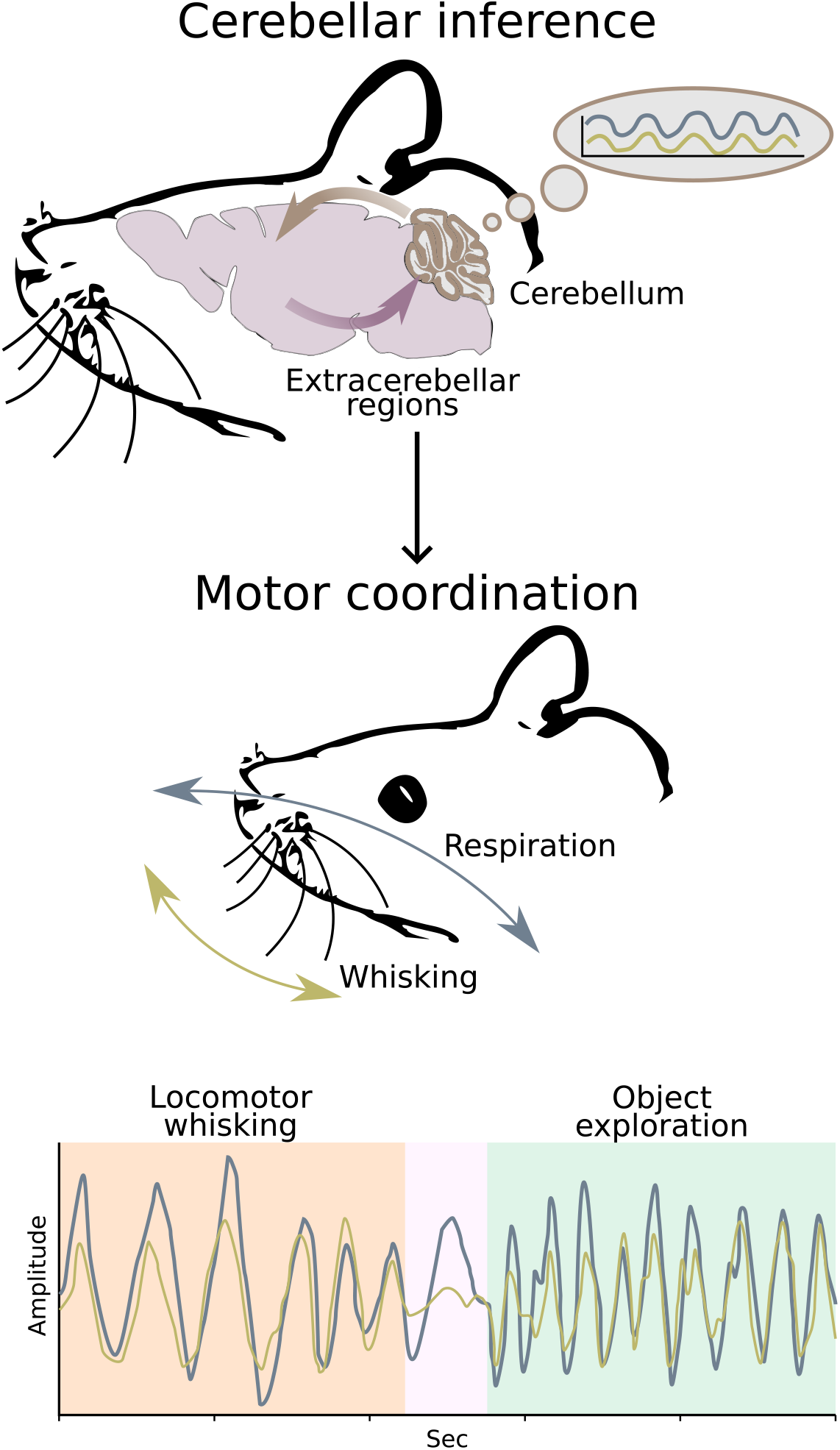
Cerebellar contribution to motor coordination. The cerebellum receives information from extra-cerebellar regions while providing top-down (prior) predictions (violet-brown cycle). Because the cerebellum has learned to expect coordinated (e.g. synchronised) dynamics between behavioural variables, such as respiration (grey line) and whisking (gold line), its predictions can contextualise behavioural domains on each others, by biasing or constraining extra-cerebellar inference. This contextualisation can span different behavioural regimes, here exemplified by a period of locomotor whisking, with low whisking and respiration rates (orange background), and a period of object exploration, with high whisking and respiration rates (green background), separated by a brief period with no whisking (pink background). Ultimately, extra-cerebellar-cerebellar interactions will therefore engender coordination between respiration and whisking.

In simulating active inference in this setting, we specify both a generative model, implemented by the cerebellum, and a generative process, the extra-cerebellar regions associated with the production of whisking and respiratory patterns. Neuronal dynamics in these regions also subtend an inference process, whereby inference is about latent states related to the whisking and inspiration-expiration cycle (e.g. their amplitude and phase); however, in this work we do not explicitly model motor behaviour, but approximate it with a stochastic process, assuming that extra-cerebellar processing engages the motor hierarchy to generate proprioceptive predictions that are realised at the level of motor reflex arcs ***Friston (2011***). Thus, by acting on this stochastic process, the cerebellum implicitly tunes the execution of whisking and respiratory behaviour (Figure 3).

In practice, we simulate whisking and respiration cycles with two Kuramoto systems, whose output *w* and *r* represent somatic states, which could be associated with whisker position and expansion of the rib cage, respectively – and are ultimately actuated through motor reflex arcs. In the Kurmoto models the two somatic states are defined by phases

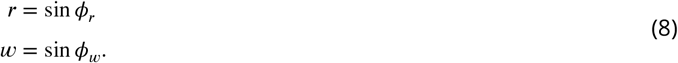

The dynamics of these phases is a function of intrinsic angular velocities, *ω*_*w*_ and *ω*_*r*_, and is subject to independent Gaussian noise *Q*_*w*_ and *Q*_*r*_ and offset *ω*_0_. Moreover, we introduce a term *α* that, together with *ω*_*w*_ and *ω*_*r*_, is state dependent, that is, it can be changed throughout simulation time, so that the *in silico* mouse can stop or restart whisking, by setting *α* to 0 or 1, or change the frequency of whisking and respiration, by modifying *ω*_*w*_ and *ω*_*r*_.

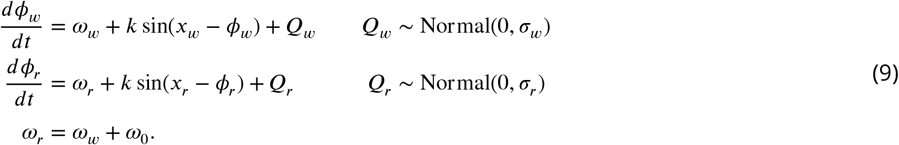

The key component of a Kuramoto system is the coupling term; in the standard Kuramoto formulation this is the sine of phase differences, and mediates the synchronisation of the oscillations. Here, in contrast, the coupling terms, *k* sin(*x*_*w*_ − *ϕ*_*w*_) and *k* sin(*x*_*r*_ − *ϕ*_*r*_) in the whisker and respiration model, relate the phases to *x*_*w*_ and *x*_*r*_, respectively. These are the corresponding hidden states inferred by the cerebellar generative model, representing extra-cerebellar somatic states *w* and *r*. The crucial point is that there is no explicit coupling between the two oscillators, instead this coupling emerges from cerebellar inferences that rest on learned expectations about synchrony. In particular, the generative model is the same state space model in (4):

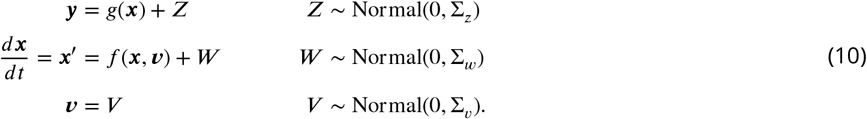

where ***y, x*** and ***v*** are now associated with whisking and respiration somatic states:

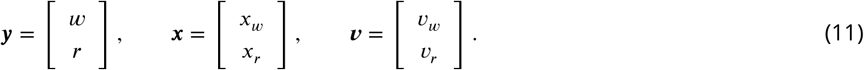

This formulation assumes that observations ***y*** – peripheral input originating from the same somatic states *w* and *r* returned by the generative process – are mapped linearly from *x*_*w*_ and *x*_*r*_, which are in turn endowed with dynamics driven by hidden causes *v*_*w*_ and *v*_*r*_. Notably, cerebellar expectations about *w*-*r* synchrony can be encoded in the **pf** connectivity *θ*_*f*_ :

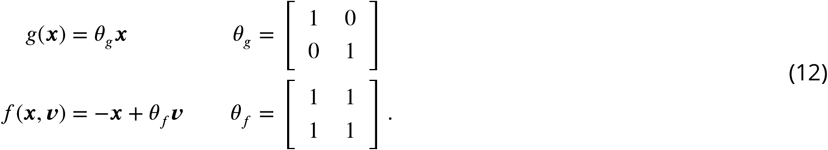

These three sets of equations (Eq. 10, Eq. 11 and Eq. 12) define the generative model, whose inversion dynamics (Eq. 6 and Eq. 7) we associate with neuronal dynamics in the cerebellum. Cerebellar state estimation is about the same whisker and respiratory states that extra-cerebellar models – here approximated by the generative process – are inferring; namely, hidden states *x*_*w*_ and *x*_*r*_ represent *w* and *r*. However, the underlying models are different: in the case of the cerebellum, the model is linear and relatively simple, conforming with empirical findings of linear encodings of many task-related parameters ***Chen et al. (2016***); ***Hong et al. (2016***); ***Raymond and Medina (2018***), whereas in the case of extra-cerebellar models, these can be nonlinear and arbitrarily complex, subtending for example the generation of cyclic patterns and nested sequences ***Kiebel and Friston (2011***); ***Kiebel et al. (2009***); ***Friston and Frith (2015***).

As such, the equations of motion governing the dynamics of somatic states in extra-cerebellar structures are synthesized in the cerebellum by a linear control of hidden states dynamics 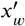 and 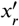 from hidden causes or control states *v*_*w*_ and *v*_*r*_. Notably, this is where the interaction between whisking and respiration occurs, as estimates of *v*_*w*_ and *v*_*r*_ (**Pjc** activity) bias through their predictions, *f* (***x, v***), estimates of *x*_*w*_ and *x*_*r*_ – replicated in **CN** activity – which in turn are sent to the generative process where they enter the coupling terms *k* sin(*x*_*w*_ − *ϕ*_*w*_) and *k* sin(*x*_*r*_ − *ϕ*_*r*_), thus forcing synchrony between the two oscillators.

Having described the setup for modelling motor coordination, we now report the results of the simulations (Figure 4, Figure 5 and Figure 6). In the following, whisking and respiration undergo state-dependent behavioural changes, which exemplify cerebellar-dependent motor coordination in different behavioural regimes. These states include a period of locomotor whisking, characterised by low rates of whisking and respiration; a brief intermediate period during which whisking stops; and a final period of object exploration, characterised by high rates of whisking and respiration (i.e., sniffing).

**Figure 4.**
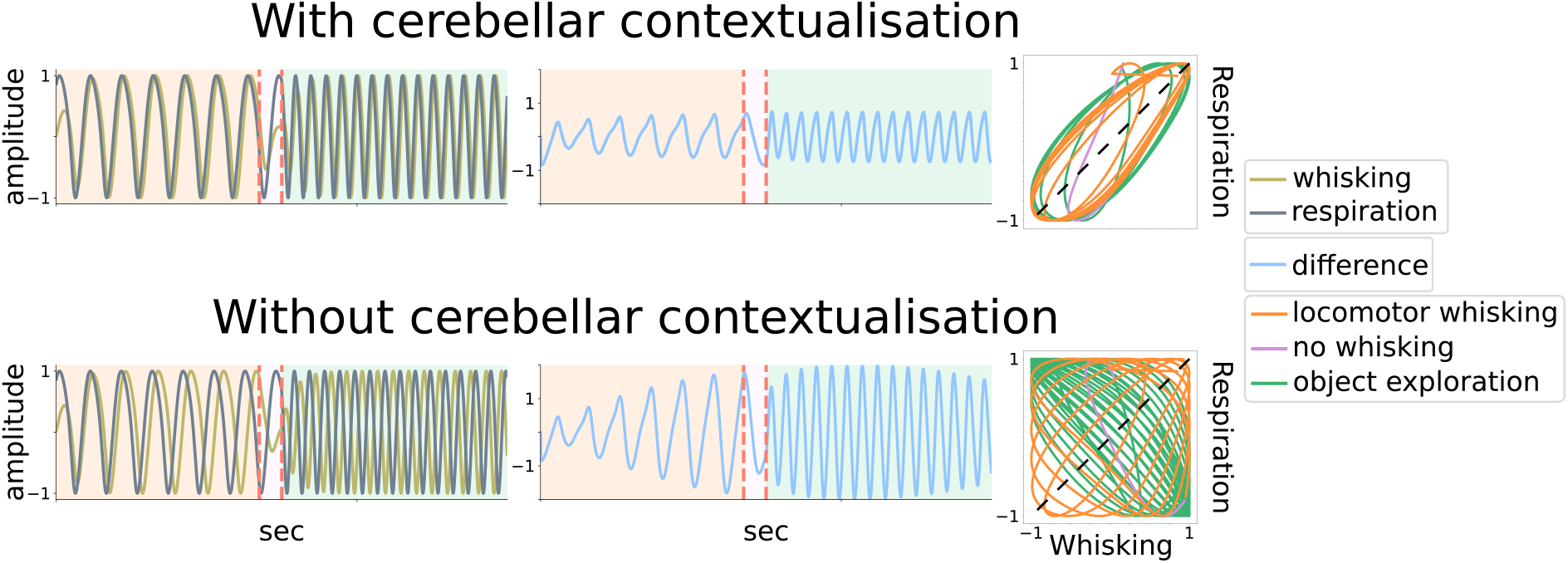
Cerebellar contribution to whisking-respiration coordination (1). Top half (‘With cerebellar contextualisation’): the cerebellar model expects coordination between whisking and respiration; these expectations in turn bias cerebellar estimates of whisking and respiration (Figure 1 - supplement 1). The left panel shows the evolution of whisking and respiration over time; the middle panel shows the evolution of their difference; the right panel shows whisking and respiration dynamics in the joint state space together with the synchronisation manifold (straight line). Behavioural state-dependent modes are color-coded, with orange, pink and green background colors (left and middle panel) or lines (right panel) corresponding to low, high whisking/respiration rate and no whisking, respectively; the transition from one mode to another is marked by vertical dotted red lines. In this simulation condition, whisking and respiration have different intrinsic angular velocities; nonetheless, in the presence of cerebellar contextualisation of extra-cerebellar dynamics, the two oscillators are kept synchronised. Bottom half (‘Without cerebellar contextualisation’): the cerebellum does not expect whisker-respiration coordination, therefore it does not contribute to the evolution of whisking and respiration. As a consequence, the two oscillators drift away from each other.

**Figure 5.**
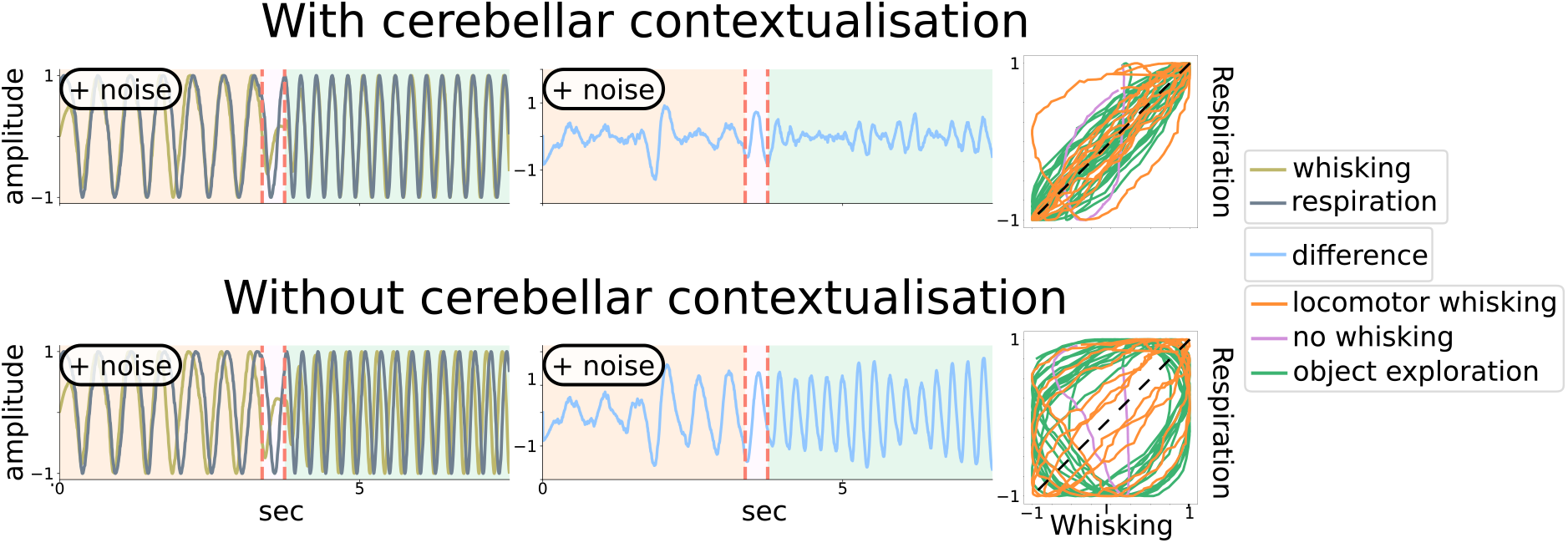
Cerebellar contribution to whisking-respiration coordination (2). Same layout as for Figure 4, but here extra-cerebellar dynamics are subject to intrinsic, independent noise. When the cerebellum expects synchrony between whisking and respiration, it is able to maintain coordination among extra-cerebellar processing; when the cerebellum does not expect synchrony, the two behaviours evolve independently. **Figure 5—video 1**. Dynamical visualisation of whisking and respiration (top) and their difference in the time domain (middle) and in the joint state space (bottom). The cerebellum expects coordination between whisking and respiration, and is able to coordinate the two behaviours despite the presence of independent gaussian noise added to their dynamics and an offset in their angular velocities (as in Figure 4). During an intermediate period, however, we simulate transient inactivation of the **CN** output to extra-cerebellar regions; consequently, whisking and respiration desynchronise. Blue and red colors indicate periods when the cerebellum does and does not couple back to extra-cerebellar structures, respectively.

**Figure 6.**
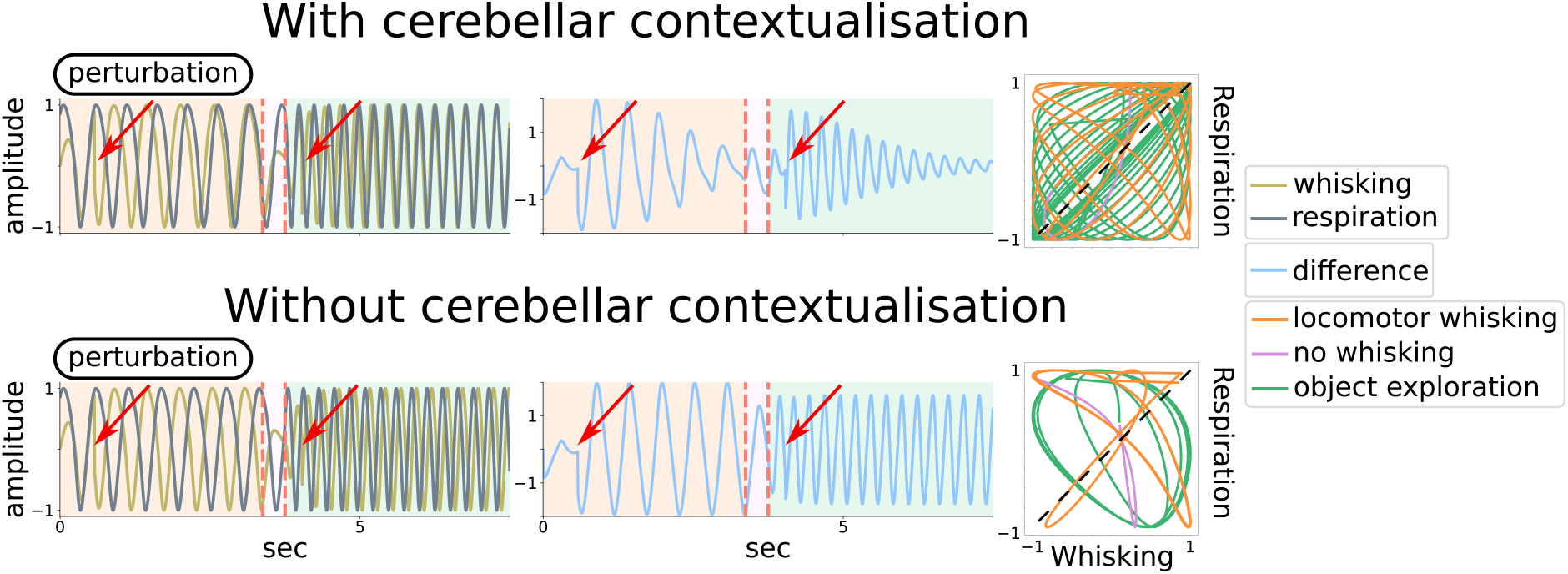
Cerebellar contribution to whisking-respiration coordination (3). Same layout as for Figure 4 and Figure 5, but here external perturbations are applied to whisking dynamics (the timings of perturbations are indicated by arrows). With cerebellar contextualisation of behaviour in play, the two oscillators return synchronised following perturbations; whereas this does not occur in the absence of cerebellar contextualisation.

The top half of Figure 4 (‘With cerebellar contextualisation’) displays the output of the generative process when the cerebellar model expects whisking-respiration coordination, that is, *θ*_*f*_ is an all-one matrix as in Eq. (11). Starting from the left panel, we see the evolution of whisker and respiratory somatic states *w* and *r*: the output of the generative process; specifically, their amplitude or displacement from a reference or resting point.

In this simulation condition, *w* and *r* have different angular velocities *ω*_*w*_ and *ω*_*r*_; this offset *ω*_0_ ≠ 0 may be due to central pattern generators having distinct autonomous dynamics, and is proposed to be partially compensated by cerebellar constraints. This is highlighted both in the middle panel, showing that the difference between *w* and *r* remains constantly below 1 a.u. (less than half the maximum difference), and in the right panel, where the two variables tend to converge onto the synchronisation manifold (dotted black line) in the joint state space.

In contrast, the bottom half of Figure 4 (‘Without cerebellar contextualisation’) shows simulation results in absence of any cerebellar expectations about behavioural coordination, that is, *θ*_*f*_ is an identity matrix. In this case, the two oscillators evolve without any cerebellar-dependent coupling. As a consequence, the difference between *w* and *r* recursively reaches its peak at 2 a.u., as the two variables are consistently out of phase, circling around the joint state space, far apart from a synchronisation manifold. Please also notice how the whisking-respiration decoupling occurs more rapidly during high compared to low oscillatory rates, which hints at an increased reliance on cerebellar computations the finer and more demanding motor control becomes.

In summary, these simulations illustrate how synchronisation of whisking and respiration is dependent on the cerebellar generative model: when this model holds expectations about motor coordination, it is able to orchestrate extracerebellar dynamics during various behavioural regimes (e.g., locomotor whisking and object exploration), despite the presence of different intrinsic rhythms and time constants. On the other hand, this cerebellar contribution is absent in the case of a naive model, namely, before learning or after lesion of **pf**.

Finally, we consider other two simulation conditions (Figure 5 and Figure 6): first, *w* and *r* dynamics are subject to independent Gaussian noise Normal(0, *σ*_w_) and Normal(0, *σ*_r_) (Figure 5); second, a perturbation occurs that abruptly changes whisking phase *ϕ*_*w*_, due to, for example, contact with an external object (Figure 6). These conditions exemplify an internal and external source of divergence between network dynamics, respectively, that the cerebellum can counteract efficiently. Accordingly, whisking and respiration are synchronised with each other in the presence of contextual modulation from the cerebellum, whereas their dynamics drift away when this contextualisation is absent.

## Discussion

We have described how state estimation, based on cerebellar internal models, can contextualise motor coordination, and more generally any aspect of behaviour that involves interactions or associations among sensorimotor states. The basic idea underlying this work is that functional behaviour is built upon state coordination and interactions, and that the cerebellum might play a key role in learning and implementing them. This is supported by empirical findings highlighting the necessity of an intact cerebellum for seamless, coordinated behaviour, resilient to perturbations ***Machado et al. (2015***, 2020). We have used – as an example – whisking-respiration coordination, where cerebellar estimates of control states underwriting synchrony are realised by modulating the activity of extra-cerebellar regions controlling whisking and respiratory patterns, respectively.

This account of cerebellar computations follows from the observations that the cerebellum receives, integrates and reciprocates concomitant information from most brain areas ***Snider and Stowell (1944***); ***Sobel et al. (1998***); ***Huang et al. (2013***); ***Chabrol et al. (2015***); ***Ishikawa et al. (2015***); ***Wagner and Luo (2020***), and its modular and relatively homogeneous architecture seems optimised to rapidly learn and infer how sensorimotor states, such as whisking and respiration, are associated or interact with each other ***Marr (1969***); ***Albus (1971***); ***Ito (2006***). As a consequence, cerebellar inference – incorporating learned relationships in its estimates – can efficiently contextualise inference in extra-cerebellar structures by providing them with timed state estimates (e.g. about whisking) that are conditioned on other states (e.g. respiration), and vice versa (Figure 1). This contextual coordination might provide a crucial advantage in coordinating behaviour, compared to coordination arising from extra-cerebellar brain regions alone, whose reciprocal communication might be subject to external conduction delays and computationally expensive.

Practically, cerebellar state estimation and the underlying neuronal dynamics comply with the FEP, that can be applied to the dynamics of biological systems from first physics principles (e.g., a principle of stationary action) ***Friston (2010***); ***Friston et al. (2022***). Under the FEP, neuronal dynamics evolve to minimise variational free energy, and neuronal networks implicitly invert probabilistic (generative or forward) models of their environment. In other words, the physical dynamics of neuronal (internal) states are equipped with belief dynamics in the space of probability distributions over (external) states. This licenses a formal link between cerebellar networks ***Apps and Hawkes (2009***) and their internal models ***Wolpert et al. (1998***); ***Ito (2006***), while providing the tools to simulate cerebellar inference. In particular, we have associated cerebellar computations with Bayesian filtering based on a state space model, which is in line with prominent models grounded on evidence of the cerebellum operating as state estimator or Kalman filter ***Paulin (1993***); ***Miall et al. (1993***); ***Tanaka et al. (2019***).

The present model – expressing average connectivity patterns and activity of neuronal populations (Figure 2) – captures key aspects of cerebellar dynamics and circuitry. First, it describes variables in terms of their trajectory in state space instead of their instantaneous value, with arbitrary accuracy; although here (for simplicity) we only consider velocity. This means that neuronal (e.g. **grc** and **Pjc**) activity can encode state trajectories, which explains the coexistance of past and future information in cerebellar activity ***Popa et al. (2012***), and affords high temporal accuracy to cerebellar state estimation.

Second, the model is linear, conforming with empirical findings of linear encoding of many task-related variables ***Chen et al. (2016***); ***Hong et al. (2016***); ***Raymond and Medina (2018***), but nonlinearities could be easily introduced in a biologically plausible manner if necessary ***Bogacz (2017***), as when dealing with delays in classical conditioning paradigms ***Thompson and Steinmetz (2009***); ***Fiocchi et al. (2022***).

Third, because we associated **Pjc** activity with the encoding of hidden causes *v*, which play the role of control states for different modes of behaviour (e.g. synchronous whisking and respiration), their activity should reflect convergent sensory and motor variables, even at a single cell level ***Gaffield et al. (2019***); ***Romano et al. (2020***).

Fourth, despite the simplicity of the model, its inversion dynamics (Eq. 6) can account for recurrency in the cerebellar architecture, involving feedback loops from **Pjc** and **CN** back to **grc, Goc** and other interneurons ***Houck and Person (2015***); ***Ankri et al. (2015***); ***Witter et al. (2016***); ***Guo et al. (2016***).

Lastly, the model is probabilistic, and takes into account uncertainty (or conversely precision) of information, which is a crucial factor for inference processes throughout the brain ***Feldman and Friston (2010***). For example, tuning the precision of extra-cerebellar input via **Goc** activity (Π_z_ in Eq. 6) may play a key role in balancing state estimation in the cerebellar cortex, and might be carefully modulated through distinct neuronal mechanisms (e.g. neuromodulatory systems) ***Palacios et al. (2021***); in the present context, the importance of changing precision parameters is evident in the simulations, with marked effects on cerebellar state estimation.

On the other hand, our simulation and model could be extended in many significant ways. In particular, the simulation of motor coordination is intentionally minimal, involving two idealised pattern generators; we could have included further somatic states, such as locomotion, head movements or licking, which have also been observed to be coordinated in rodents ***Kurnikova et al. (2017***); ***McElvain et al. (2018***); ***Grant et al. (2018***). Furthermore, it would be interesting to apply the same cerebellar model to different task conditions, such as perceptual processing of behaviourally relevant states ***Baumann et al. (2015***) or decision making ***Gao et al. (2018***); ***Deverett et al. (2018***); ***Sendhilnathan et al. (2020***).

As for the model itself, a first crucial addition would include the olivary-climbing fibre system next to the **mf** pathway. Climbing fibres (**cf**) carry many different types of sensory, motor or cognitive input ***Fu et al. (1997***); ***Deverett et al. (2018***); ***Ju et al. (2019***); ***Heffley and Hull (2019***); ***Ikezoe et al. (2022***), reflecting again the global involvement of the cerebellum in behaviour. Importantly, **cf** input can be treated similarly to **pf** input, namely, as prediction errors, driving (Bayesian) belief updating in **Pjc**; interestingly, these updates should inform **Pjc** about (hidden) states that are particularly informative of behavioural contingencies (e.g. unconditioned stimulus in conditioning paradigms), in the sense that they would entrain learning interactions or associations between other behaviourally relevant states (e.g. a conditioned stimulus with unconditioned responses), in the form of changes in the pf connectivity, 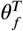 (see for example ***Friston and Herreros (2016***)).

A second important extension to the model would involve a more detailed characterisation of **CN**. In particular, we have assumed that they receive the same **mf** input as the cerebellar cortex, and thus modelled them by simply replicating hidden states *x* encoded in **grc**; however, this is not necessarily the case ***Tanaka et al. (2020***), which would motivate the addition of different hidden states encoded in **CN** and complementing those in **grc**.

Finally, it is interesting to consider the application of the cerebellar model presented here when simulating inference throughout the brain. A first point concerns the modularity of the extra-cerebellar generative model (e.g. visual and auditory pathways), which naturally emerges from conditional independencies among variables ***Parr et al. (2020***). Mathematically, interactions between modules can be mediated by mean field terms, meaning that only the average estimate of one module influences estimates of another module; in this case, the cerebellum might provide efficient estimates of control states that are conditioned on other variables.

A second point pertains to the method used to simulate cerebellar inference: in doing so, one could either simulate full cerebellar state estimation – using inversion dynamics based on a model of how *x* generates *y* – or approximate the latter using a recognition model, namely, a function mapping observations *y* to states *x* (Figure 7). The implementation of a cerebellar recognition model can be achieved with an adaptive filter ***Fujita (1982***); ***Porrill et al. (2013***) or, more generally, with a neural network, whose internal dynamics is equivalent to variational Bayesian inference used here to model state estimation ***Isomura (2022***). In practice, this would allow one to approximate a simple cerebellar neuronal inference scheme with an even simpler and computationally more efficient mapping, that is, one could provide extracerebellar structures with estimates of hidden states *x* – contextualised by other *x*’s inferred in other brain regions – that are directly mapped from observations *y*’s. However, this would confound the clear link between between cerebellar computations (state estimation) supporting coordinated behaviour and its neuronal dynamics described in this work.

**Figure 7.**
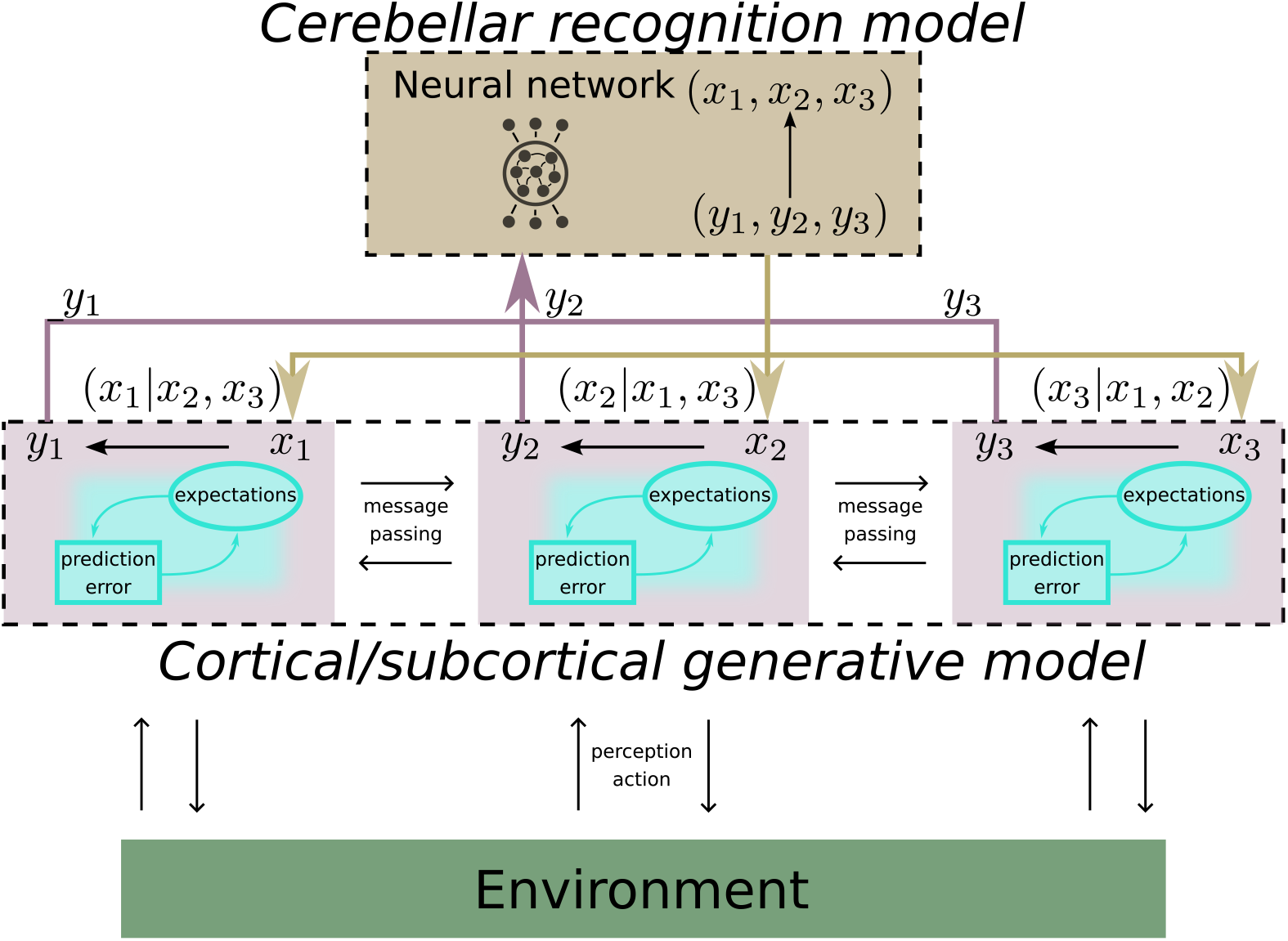
Cerebellar (recognition) model contribution to neuronal inference in the brain. This schematic is similar to Figure 1, but here the cerebellar generative model – describing how states *x* produce observations *y* – is approximated by a recognition model, directly mapping *y* to *x*, implemented by a neural network whose dynamics are equivalent to variational Bayesian inference used in our simulation of cerebellar state estimates ***Isomura (2022***). This substitution has the advantage of simplifying computations, as now there is no need to simulate the full inversion dynamics based on the original cerebellar model, but at the expense of direct link between cerebellar computations (state estimation) and its neuronal dynamics. Notably, the function implemented by the neural network also accounts for the fact that different *y*′s contextualise each others when mapping onto *x*′s, similarly to the cerebellar generative model in Figure 1 when learning interactions between *x*′s in generating *y*.

## Conclusion

We have presented a minimal cerebellar model for state estimation of sensorimotor states, and described how it could underwrite motor coordination, and more generally contextualise and finesse behaviour. Briefly, the key aspect of this contribution relies on the fact that the cerebellum is optimised to learn how different behavioural states are – or should be – coordinated with each other, and efficiently realise these learned expectations by biasing inference in extra-cerebellar brain regions accordingly. The model complies with active inference, which affords a formal link between neuronal dynamics in the cerebellum and probabilistic (generative or forward) models entailed. To substantiate our claims, we have shown how the components of such models may map onto elements of the cerebellar architecture, and provided simulations of cerebellar state estimation in the context of whisking-respiration coordination in mouse as proof of concept. Finally, we have discussed how our model accounts for several important aspects of cerebellar computations and dynamics, how it could be expanded, and how it could be efficiently included in simulations of whole-brain neuronal inference.

## Methods and Materials

### Variational Free energy

In Bayesian inference, we seek the posterior distribution over unknown states *ϑ*, given some observations *y* and a model *m* of how these observations have been generated:

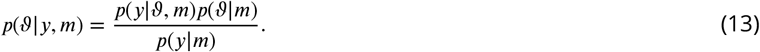

Finding the posterior is generally intractable due to the calculation of the marginal likelihood or normalisation constant (a.k.a. model evidence) *p*(*ϑ* |*m*). Therefore it has been suggested that the brain approximates Bayesian inference using variational Bayes, an optimisation procedure that allows us to find a variational or recognition density over environmental states, *q*(*ϑ*), which approximates the true posterior *p*(*ϑ*|*y*). This optimisation relies on the minimisation of the Kullback-Liebler (KL) divergence between the proposed and true posterior density. Importantly, the KL divergence can be expressed as a variational free energy (F) ***Buckley et al. (2017***); leaving out the *m* for clarity, we substitute from the Bayes law, Equation 13

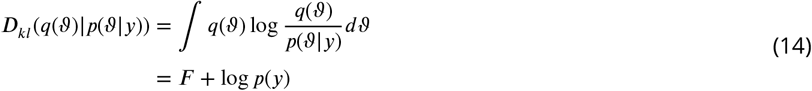

where

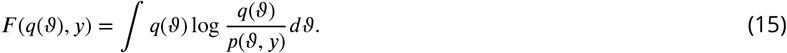

*F* depends on the generative model *m*, since *p*(*ϑ, y*), as does the recognition density *q*(*ϑ*). The sufficient statistics (e.g., mean or expectation) of *q*(*ϑ*) are assumed to be encoded by the brain. The equation for the KL divergence, Equation 14, shows that minimising *F* by changing *q*(*ϑ*) automatically reduces *D*_*kl*_, because log *p*(*y*) does not depend on *q*(*ϑ*). Thus, minimising the free energy *F* makes the recognition density, *q*(*ϑ*) a good proxy for *p*(*ϑ*|*y*).

This leaves open the question of what form *q*(*ϑ*) takes; under the Laplace approximation it is restricted to a Gaussian distribution. Thus, in this picture, the brain encodes *q*(*ϑ*) through the sufficient statistics *μ* and *ξ*, the mean and variance respectively, of a Gaussian distribution *q*(*ϑ*; *μ, ξ*) ***Friston et al. (2008***), effectively constraining beliefs about hidden or latent states to be normally distributed. Under the Laplace assumption, the variance of the recognition density that minimises free energy is an analytic function of the expectation. This means that the brain only needs to encode the expected value *μ* of these beliefs in order to minimise *F*. The functional form of *F* is specified by *m*, which is, in this work, a state space model with first-order generalised motion (bold face refers to vectors) ***Friston et al. (2008***):

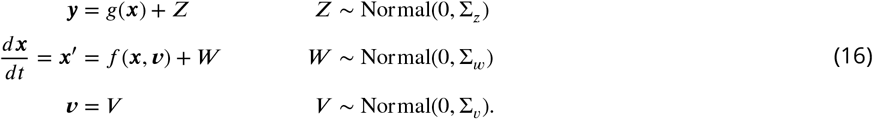

This model assumes the existence of two hidden or latent states, ***v*** and ***x***, which bring about observations ***y***; ***v*** can be thought of as the control or exogenous input to a system and ***x*** as the state of a system whose temporal dynamics ***x***′ are a function of itself and the control states ***v***. On the other hand, *Z* and *W* are noise terms affecting the mapping *g* from ***x*** to ***y*** and equations of motion for ***x***; *V* is a noise term with high variance, indicating a noninformative prior over ***v***. With this model, the free energy can then be decomposed as (omitting some terms for clarity):

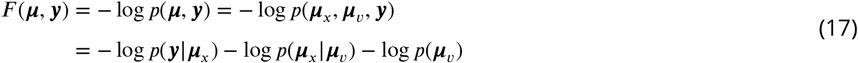

and, using the model, as described in Equation 16, these log probabilities can be written as:

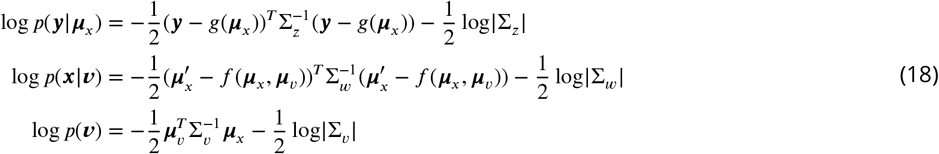

which in turn gives the following functional form:

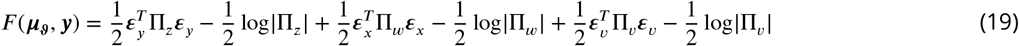

where we have used the precision matrix in place the covariance matrix (Π = Σ^−1^) and the auxiliary variables ***ε*** to indicate the prediction error terms:

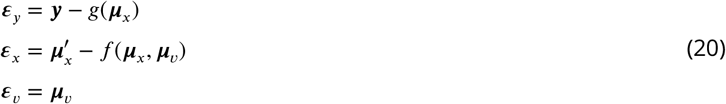

where

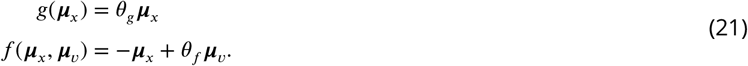

It is proposed that neuronal dynamics can be understood as minimising *F*. This can be achieved by specifying their temporal evolution as a gradient descent on *F* :

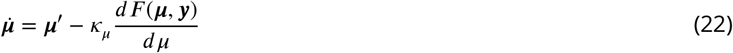

It is important to note that in this gradient descent scheme, the term updating expectations about beliefs encoded by neuronal dynamics, 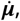 is distinct from the motion of those expectations, ***μ***^′^; that is, the motion of the expectations is distinct from expected motion. The two equate only when *F* is minimised ***Friston et al. (2008***). The equation of motion, Equation (22), describes the general form of *recognition dynamics* underlying perception, that is, the dynamics of neuronally encoded states which minimise *F*. Alternatively, minimisation of *F* can be achieved by changing the state of the environment through action *a*, so that observations conform to beliefs ***Friston (2011***); ***Friston et al. (2016***). Action dynamics can be specified with a gradient descent on *F* :

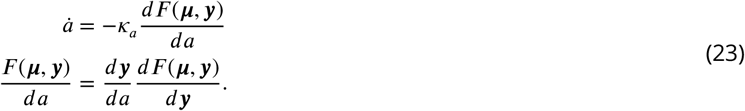

In this formulation, action *a* does not enter in the definition of *F* (***μ, y***), because the brain does not entertain any explicit belief about *a* in its model *m* about the environment. However, the brain possesses an *inverse model* of how *a* acts on the environment and changes ***y***: this could be implemented for example by a cascade of belief propagation down to the level of arc reflexes in the spinal cord, hardwired to satisfy expectations about limb position by producing the appropriate pattern of muscle contractions ***Friston (2011***). In the present work, because of the way we couple the cerebellum to extracerebellar structures through estimates of *x*, we do not deal with *a* explicitly. As such, we only focus on the recognition dynamics for ***x, x***′, ***v*** based on the internal energy, Equation (19). This gives dynamical equations for the means:

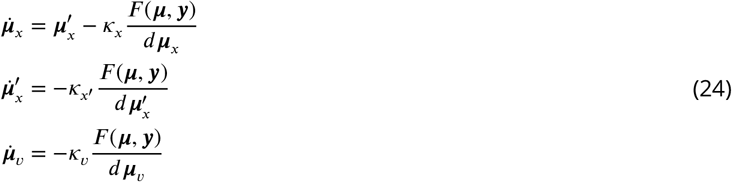

Here the expected motion 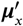 only enters the recognition dynamics of ***μ***_***x***_, because for 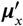 there is no expected motion, that is, there is no s 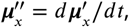, and *v* does not have any dynamics. Finally, the partial derivatives in equations of motions for the means, Equation 24, can be expanded into equations for the rate of change of *F* (***μ, y***) with respect to ***μ***_***x***_:

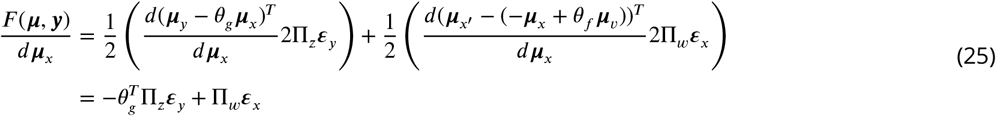

with respect to ***μ***_*x*′_ :

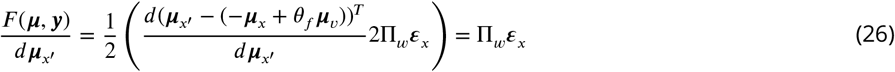

and with respect to ***μ***_***v***_:

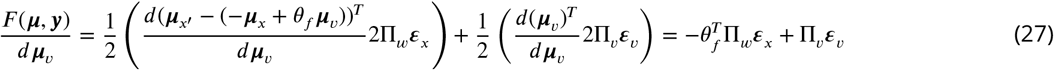

where we used the matrix differentiation property

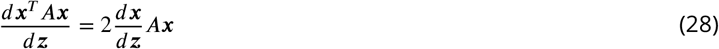

for symmmetric *A*.

### FEP narrative summary

The free energy principle (FEP) is a theory of self-organisation ***Friston (2010***), purported to explain – from first principles – the existence and persistence of a system embedded in its environment. A key result of the FEP is the interpretation of the dynamics of a self-organising system (like the brain) as emulating an inference process ***Ramstead et al. (2022***). A narrative summary goes as follows: the FEP starts from the observation of what it means for a system to exist, namely, to be distinguishable from, but in exchange with, its environment; this basic intuition is formalised by appealing to the presence of a statistical boundary, a Markov blanket, both instantiating a separation and mediating the exchange between internal states of the system and states of the environment ***Kirchhoff et al. (2018***); ***Palacios et al. (2020***). Alongside this ontological description of a system – as possessing a Markov blanket – its existence induces a probability density over states that are characteristic of the system in question. This density places constraints on dynamics or trajectories in state space of internal and active blanket states (known as autonomous states ***Friston (2013***); ***Karl et al. (2022***). From here, FEP proceeds to express the dynamics of autonomous states as a principle of least action: ^1^ intuitively, this principle prescribes changes in the states of the system, subject to the boundary condition that their trajectories minimise a certain quantity, called the Lagrangian. Under the FEP, this quantity equates to the aforementioned (negative log) probability over trajectories, such that the dynamics of the system will, on average, maximise this density, that is, to conform to the most likely paths; those characterising the system in the first instance. Notably, the initial description of the system based on its Markov blanket underpins a stipulative mapping between the internal states and a probability distribution over external states, conditioned on the blanket states; this mapping is central to the FEP, because it couples the physical description of a system with a description of it in terms of a probabilistic model, parametrised by internal states, about its environment.

Mathematically, this enables one to express internal and blanket trajectories, as a principle of least action, that minimises the path integral of free energy (F) functional that shares the same minima as the Lagrangian ***Ramstead et al. (2022***). This variational free energy is used in approximate Bayesian inference to score the goodness of a probabilistic (i.e., generative) model of the environment, given some observations. Consequently, the most likely trajectories of the system’s states are those that comply with its ontological description and at the same time maximise the evidence of an implicit model of the environment encoded by internal states of the system. Ultimately, this formalisation of self-organisation is what licences an interpretation of internal dynamics as an inference process.

---

**Figure 4—figure supplement 1.**
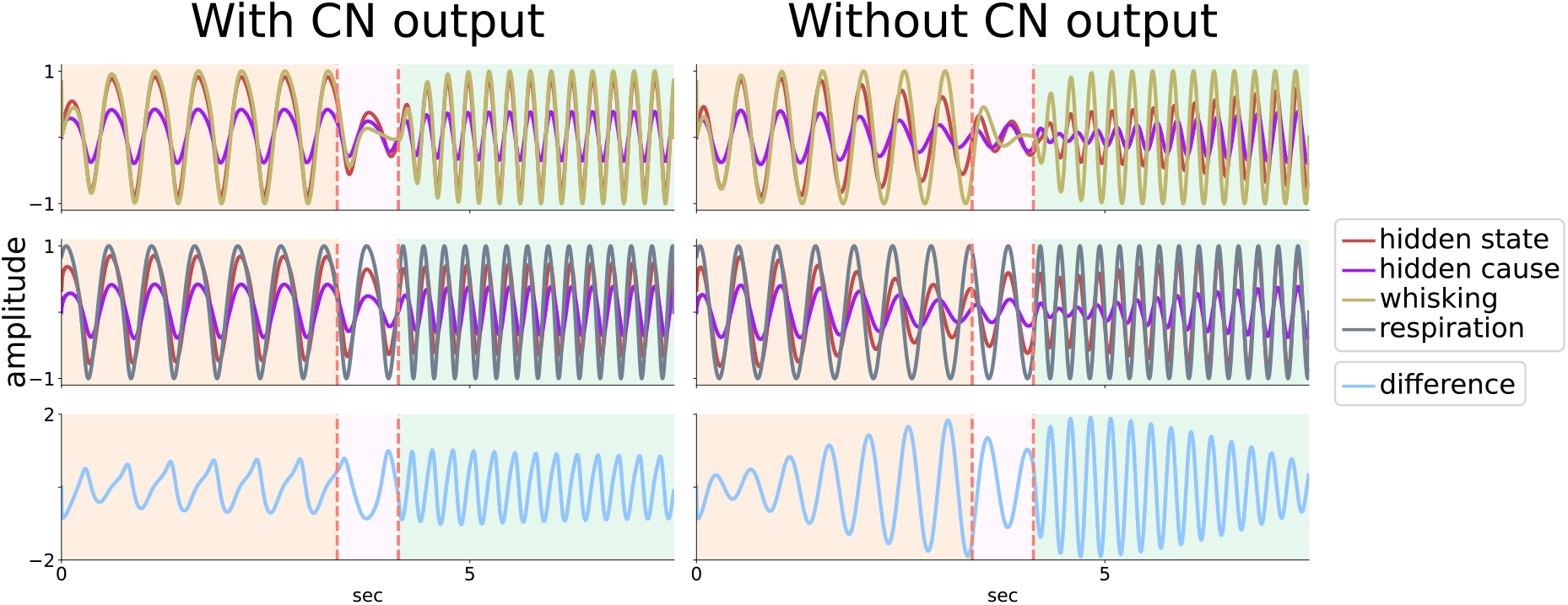
Cerebellar expectations bias state estimation. Simulation results in which the cerebellum expects synchrony between whisking and respiration, whose intrinsic dynamics have different angular velocities. Whisking and respiration variables are split in the top and middle panels, respectively, and are displayed together with cerebellar estimates of the associated hidden states and causes. The bottom panel displays their difference. Left: the cerebellum sends empirical priors via **CN** outputs to extra-cerebellar structures. In this case, estimates about hidden states *x*_*w*_ and *x*_*r*_ (red lines) closely match real states *w* and *r* – in virtue of the fact that *w* and *r* are constrained towards *x*_*w*_ and *x*_*r*_ – while estimates about hidden causes *v*_*w*_ and *v*_*w*_ (violet lines) are regular and relatively high in amplitude, which reflects the congruence between expected and observed synchrony between behavioural states. Right: the output of **CN** to extra-cerebellar structures is suspended: now, estimates about hidden states and causes reduce in amplitude the more behaviour desynchronises. This is because there is a friction between predictions and observations, which has an impact on estimates about hidden causes – a function of behavioural synchrony due to expectations in *θ*_*f*_ – as well as hidden states – driven by contrasting ascending information and feedback predictions within the cerebellum.

## Footnotes

1 The principle of least action is used when dealing with probabilities of trajectories over states, as opposed to dynamics of probabilities over states, in which case the FEP uses a path integral formulation ***Ramstead et al. (2022***); ***Friston et al. (2022***)

## References

Albus JS. A theory of cerebellar function. Mathematical Biosciences. 1971; 10(1):25–61.

Ankri L, Husson Z, Pietrajtis K, Proville R, Léna C, Yarom Y, Dieudonné S, Uusisaari MY. A novel inhibitory nucleo-cortical circuit controls cerebellar Golgi cell activity. eLife. 2015; 4:e06262.

Apps R, Hawkes R. Cerebellar cortical organization: a one-map hypothesis. Nature Reviews Neuroscience. 2009; 10(9):670–681.

Argyropoulos GPD. The cerebellum, internal models and prediction in ‘non-motor’ aspects of language: A critical review. Brain and Language. 2016; 161:4–17.

Arshavsky YI, Deliagina TG, Orlovsky GN. Pattern generation. Current Opinion in Neurobiology. 1997; 7(6):781–789.

Baumann O, Borra RJ, Bower JM, Cullen KE, Habas C, Ivry RB, Leggio M, Mattingley JB, Molinari M, Moulton EA, Paulin MG, Pavlova MA, Schmahmann JD, Sokolov AA. Consensus paper: The role of the cerebellum in perceptual processes. Cerebellum. 2015; 14(2):197–220.

Baumann O, Mattingley JB. Cerebellum and emotion processing. In: Adamaszek M, Manto M, Schutter DJLG, editors. The Emotional Cerebellum Cham: Springer International Publishing; 2022. p. 25–39.

Becker MI, Person AL. Cerebellar control of reach kinematics for endpoint precision. Neuron. 2019; 103(2):335–348.e5.

Bellingham MC. Driving respiration: The respiratory central pattern generator. Clinical and Experimental Pharmacology and Physiology. 1998; 25(10):847–856.

Bergmann R, Sehara K, Dominiak SE, Kremkow J, Larkum ME, Sachdev RNS. Coordination between eye movement and whisking in head-fixed mice navigating a plus maze. eNeuro. 2022; 9(4).

Bogacz R. A tutorial on the free-energy framework for modelling perception and learning. Journal of Mathematical Psychology. 2017; 76:198 –211.

Brooks JX, Carriot J, Cullen KE. Learning to expect the unexpected: rapid updating in primate cerebellum during voluntary self-motion. Nature Neuroscience. 2015; 18(9):1310–1317.

Buckley CL, Kim CS, McGregor S, Seth AK. The free energy principle for action and perception: A mathematical review. Journal of Mathematical Psychology. 2017; 81:55–79.

Cerminara NL, Apps R, Marple-Horvat DE. An internal model of a moving visual target in the lateral cerebellum. Journal of Physiology. 2009; 587(2):429–442.

Chabrol FP, Arenz A, Wiechert MT, Margrie TW, Digregorio DA. Synaptic diversity enables temporal coding of coincident multisensory inputs in single neurons. Nature Neuroscience. 2015; 18(5):718–727.

Chen S, Augustine GJ, Chadderton P. The cerebellum linearly encodes whisker position during voluntary movement. eLife. 2016; 5.

Coesmans M, Weber JT, De Zeeuw CI, Hansel C. Bidirectional parallel fiber plasticity in the cerebellum under climbing fiber control. Neuron. 2004; 44(4):691–700.

Da Costa L, Parr T, Sengupta B, Friston K. Neural dynamics under active inference: Plausibility and efficiency of information processing. Entropy. 2021; 23(4).

Dacre J, Colligan M, Clarke T, Ammer JJ, Schiemann J, Chamosa-Pino V, Claudi F, Harston JA, Eleftheriou C, Pakan JMP, Huang CC, Hantman AW, Rochefort NL, Duguid I. A cerebellar-thalamocortical pathway drives behavioral context-dependent movement initiation. Neuron. 2021; 109(14):2326–2338.e8.

D’Angelo E. The organization of plasticity in the cerebellar cortex: From synapses to control. In: Ramnani N, editor. Cerebellar Learning, vol. 210 of Progress in Brain Research Elsevier; 2014.p. 31–58.

Deverett B, Koay SA, Oostland M, Wang SSH. Cerebellar involvement in an evidence-accumulation decision-making task. eLife. 2018; 7:e36781.

Doya K. What are the computations of the cerebellum, the basal ganglia and the cerebral cortex? Neural Networks. 1999; 12(7):961–974.

Feldman AG, Levin MF. The origin and use of positional frames of reference in motor control. Behavioral and Brain Sciences. 1995; 18(4):723–744.

Feldman H, Friston K. Attention, uncertainty, and free-energy. Frontiers in Human Neuroscience. 2010; 4:215.

Fiocchi FR, Dijkhuizen S, Koekkoek SKE, De Zeeuw CI, Boele HJ. Stimulus generalization in mice during Pavlovian eyeblink conditioning. eNeuro. 2022; 9(2).

Fonio E, Gordon G, Barak N, Winetraub Y, Oram TB, Haidarliu S, Kimchi T, Ahissar E. Coordination of sniffing and whisking depends on the mode of interaction with the environment. Israel Journal of Ecology and Evolution. 2015; 61(2):95–105.

Frens M, Donchin O. Forward models and state estimation in compensatory eye movements. Frontiers in Cellular Neuroscience. 2009;

Friston K. The free-energy principle: a rough guide to the brain? Trends in Cognitive Sciences. 2009; 13(7):293–301.

Friston K. The free-energy principle: a unified brain theory? Nature Reviews Neuroscience. 2010; 11(2):127–138.

Friston K. What is optimal about motor control? Neuron. 2011; 72(3):488–498.

Friston K. Life as we know it. Journal of The Royal Society Interface. 2013; 10(86):20130475.

Friston K, Da Costa L, Sajid N, Heins C, Ueltzhöffer K, Pavliotis GA, Parr T, The free energy principle made simpler but not too simple. arXiv; 2022.

Friston K, FitzGerald T, Rigoli F, Schwartenbeck P, Pezzulo G. Active inference: A process theory. Neural Computation. 2016; 29(1):1–49.

Friston K, Frith C. A duet for one. Consciousness and Cognition. 2015; 36:390–405.

Friston K, Herreros I. Active inference and learning in the cerebellum. Neural Computation. 2016; 28(9):1812–1839.

Friston K, Kiebel S. Predictive coding under the free-energy principle. Philosophical Transactions of the Royal Society B: Biological Sciences. 2009; 364(1521):1211–1221.

Friston K, Mattout J, Kilner J. Action understanding and active inference. Biological Cybernetics. 2011; 104(1-2):137–160.

Friston KJ, Trujillo-Barreto N, Daunizeau J. DEM: A variational treatment of dynamic systems. NeuroImage. 2008; 41(3):849–885.

Frontera JL, Léna C. When the cerebellum holds the starting gun. Neuron. 2021; 109(14):2207–2209.

Fu QG, Mason CR, Flament D, Coltz JD, Ebner TJ. Movement kinematics encoded in complex spike discharge of primate cerebellar Purkinje cells. Neuroreport. 1997; 8(2):523–529.

Fujita M. Adaptive filter model of the cerebellum. Biological Cybernetics. 1982; 45(3):195–206.

Gaffield MA, Bonnan A, Christie JM. Conversion of graded presynaptic climbing fiber activity into graded postsynaptic Ca2+ signals by Purkinje cell Dendrites. Neuron. 2019; 102(4):762–769.e4.

Gao Z, Davis C, Thomas AM, Economo MN, Abrego AM, Svoboda K, De Zeeuw CI, Li N. A cortico-cerebellar loop for motor planning. Nature. 2018; 563(7729):113–116.

Golomb D, Moore JD, Fassihi A, Takatoh J, Prevosto V, Wang F, Kleinfeld D. Theory of hierarchically organized neuronal oscillator dynamics that mediate rodent rhythmic whisking. Neuron. 2022; 110(22):3833–3851.e22.

Grant RA, Breakell V, Prescott TJ. Whisker touch sensing guides locomotion in small, quadrupedal mammals. Proceedings of the Royal Society B: Biological Sciences. 2018; 285(1880):20180592.

Guo C, Witter L, Rudolph S, Elliott H, Ennis K, Regehr W. Purkinje cells directly inhibit granule cells in specialized regions of the cerebellar cortex. Neuron. 2016; 91(6):1330–1341.

Hansel C, Linden DJ, D’Angelo E. Beyond parallel fiber LTD: the diversity of synaptic and non-synaptic plasticity in the cerebellum. Nature Neuroscience. 2001; 4(5):467–475.

Heffley W, Hull C. Classical conditioning drives learned reward prediction signals in climbing fibers across the lateral cerebellum. Elife. 2019; 8.

Hohwy J. The self-evidencing brain. Noûs. 2016; 50(2):259–285.

Hong S, Negrello M, Junker M, Smilgin A, Thier P, De Schutter E. Multiplexed coding by cerebellar Purkinje neurons. eLife. 2016; 5:e13810.

Houck BD, Person AL. Cerebellar premotor output neurons collateralize to innervate the cerebellar cortex. Journal of Comparative Neurology. 2015; 523(15):2254–2271.

Huang CC, Sugino K, Shima Y, Guo C, Bai S, Mensh BD, Nelson SB, Hantman AW. Convergence of pontine and proprioceptive streams onto multimodal cerebellar granule cells. eLife. 2013; 2.

Ikezoe K, Hidaka N, Manita S, Murakami M, Tsutsumi S, Isomura Y, Kano M, Kitamura K. Cerebellar climbing fibers convey behavioral information of multiplex modalities and form functional modules. bioRxiv. 2022;.

Ishikawa T, Shimuta M, Häusser M. Multimodal sensory integration in single cerebellar granule cells in vivo. eLife. 2015; 4.

Isomura T. Active inference leads to Bayesian neurophysiology. Neuroscience Research. 2022; 175:38–45. Constructive Understanding of Multi-scale Dynamism of Neuropsychiatric Disorders.

Ito M. Cerebellar circuitry as a neuronal machine. Progress in Neurobiology. 2006; 78(3):272–303. The Contributions of John Carew Eccles to Contemporary Neuroscience.

Jirsa VK, Kelso JAS. The excitator as a minimal model for the coordination dynamics of discrete and rhythmic movement generation. Journal of Motor Behavior. 2005; 37(1):35–51.

Ju C, Bosman LWJ, Hoogland TM, Velauthapillai A, Murugesan P, Warnaar P, van Genderen RM, Negrello M, De Zeeuw CI. Neurons of the inferior olive respond to broad classes of sensory input while subject to homeostatic control. Journal of Physiology. 2019; 597(9):2483–2514.

Karl F, Lancelot DC, Dalton ARS, Conor H, Grigorios AP, Maxwell R, Parr T, Path integrals, particular kinds, and strange things. arXiv; 2022.

Kelly E, Meng F, Fujita H, Morgado F, Kazemi Y, Rice LC, Ren C, Escamilla CO, Gibson JM, Sajadi S, Pendry RJ, Tan T, Ellegood J, Basson MA, Blakely RD, Dindot SV, Golzio C, Hahn MK, Katsanis N, Robins DM, et al. Regulation of autism-relevant behaviors by cerebellar– prefrontal cortical circuits. Nature Neuroscience. 2020; 23(9):1102–1110.

Kelso JAS. Unifying large- and small-scale theories of coordination. Entropy. 2021; 23(5).

Kiebel SJ, Daunizeau J, Friston KJ. A hierarchy of time-scales and the brain. PLOS Computational Biology. 2008; 4(11):1–12.

Kiebel SJ, Friston KJ. Free energy and dendritic self-organization. Frontiers in Systems Neuroscience. 2011; 5:80.

Kiebel SJ, von Kriegstein K, Daunizeau J, Friston KJ. Recognizing sequences of sequences. PLOS Computational Biology. 2009; 5(8):1–13.

Kirchhoff M, Parr T, Palacios E, Friston K, Kiverstein J. The Markov blankets of life: autonomy, active inference and the free energy principle. Journal of The Royal Society Interface. 2018; 15(138):20170792.

Koziol LF, Budding D, Andreasen N, D’Arrigo S, Bulgheroni S, Imamizu H, Ito M, Manto M, Marvel C, Parker K, Pezzulo G, Ramnani N, Riva D, Schmahmann J, Vandervert L, Yamazaki T. Consensus paper: The cerebellum’s role in movement and cognition. Cerebellum. 2014; 13(1):151–177.

Kurnikova A, Moore JD, Liao SM, Deschênes M, Kleinfeld D. Coordination of orofacial motor actions into exploratory behavior by rat. Current Biology. 2017; 27(5):688–696.

Li N, Mrsic-Flogel TD. Cortico-cerebellar interactions during goal-directed behavior. Current Opinion in Neurobiology. 2020; 65:27–37.

Lin Q, Manley J, Helmreich M, Schlumm F, Li JM, Robson DN, Engert F, Schier A, Nöbauer T, Vaziri A. Cerebellar neurodynamics predict decision timing and outcome on the single-trial level. Cell. 2020; 180(3):536–551.e17.

Liu X, Robertson E, Miall RC. Neuronal activity related to the visual representation of arm movements in the lateral cerebellar cortex. Journal of Neurophysiology. 2002; 89(3):1223–1237.

Liu X, Robertson E, Miall RC. Neuronal activity related to the visual representation of arm movements in the lateral cerebellar cortex. Journal of Neurophysiology. 2003; 89(3):1223–1237.

Liu Y, Qi S, Thomas F, Correia BL, Taylor AP, Sillitoe RV, Heck DH. Loss of cerebellar function selectively affects intrinsic rhythmicity of eupneic breathing. Biology Open. 2020; 9(4). Bio048785.

Machado AS, Darmohray DM, Fayad J, Marques HG, Carey MR. A quantitative framework for whole-body coordination reveals specific deficits in freely walking ataxic mice. eLife. 2015 oct; 4:e07892.

Machado AS, Marques HG, Duarte DF, Darmohray DM, Carey MR. Shared and specific signatures of locomotor ataxia in mutant mice. eLife. 2020 jul; 9:e55356.

Mansell W. Control of perception should be operationalized as a fundamental property of the nervous system. Topics in Cognitive Science. 2011; 3(2):257–261.

Marr D. A theory of cerebellar cortex. Journal of Physiology. 1969; 202(2):437–470.

Mauk MD, Ruiz BP. Learning-dependent timing of Pavlovian eyelid responses: Differential conditioning using multiple interstimulus intervals. Behavioral Neuroscience. 1992; 106:666–681.

McElvain LE, Friedman B, Karten HJ, Svoboda K, Wang F, Deschênes M, Kleinfeld D. Circuits in the rodent brainstem that control whisking in concert with other orofacial motor actions. Neuroscience. 2018; 368:152–170. Barrel Cortex Function.

Miall RC, Weir DJ, Wolpert DM, Stein JF. Is the cerebellum a Smith predictor? Journal of Motor Behavior. 1993; 253:203–16.

Palacios ER, Houghton C, Chadderton P. Accounting for uncertainty: inhibition for neural inference in the cerebellum. Proceedings of the Royal Society B: Biological Sciences. 2021; 288(1947):20210276.

Palacios ER, Isomura T, Parr T, Friston K. The emergence of synchrony in networks of mutually inferring neurons. Scientific Reports. 2019; 9(1):6412.

Palacios ER, Razi A, Parr T, Kirchhoff M, Friston K. On Markov blankets and hierarchical self-organisation. Journal of Theoretical Biology. 2020; 486:110089.

Parr T, Sajid N, Friston KJ. Modules or Mean-Fields? Entropy. 2020; 22(5).

Paulin MG. Evolution of the cerebellum as a neuronal machine for Bayesian state estimation. Journal of Neural Engineering. 2005; 2(3):S219–S234.

Paulin MG. A model of the role of the cerebellum in tracking and controlling movements. Human Movement Science. 1993; 12(1):5–16.

Popa L, Hewitt A, Ebner T. Predictive and feedback performance errors are signaled in the simple spike discharge of individual Purkinje cells. Journal of Neuroscience. 2012; 32:15345–58.

Porrill J, Dean P, Anderson SR. Adaptive filters and internal models: Multilevel description of cerebellar function. Neural Networks. 2013; 47:134–149.

Ramnani N. Automatic and controlled processing in the corticocerebellar system. In: Ramnani N, editor. Cerebellar Learning, vol. 210 of Progress in Brain Research Elsevier; 2014.p. 255–285.

Ramstead MJD, Sakthivadivel DAR, Heins C, Koudahl M, Millidge B, Da Costa L, Klein B, Friston KJ, On Bayesian mechanics: A physics of and by beliefs. arXiv; 2022.

Raymond JL, Medina JF. Computational principles of supervised learning in the cerebellum. Annual Review of Neuroscience. 2018; 41(1):233–253.

Romano V, Reddington AL, Cazzanelli S, Mazza R, Ma Y, Strydis C, Negrello M, Bosman LWJ, De Zeeuw CI. Functional convergence of autonomic and sensorimotor processing in the lateral cerebellum. Cell Reports. 2020; 32(1):107867.

Rondi-Reig L, Paradis AL, Fallahnezhad M. A liaison brought to light: Cerebellum-Hippocampus, partners for spatial cognition. Cerebellum. 2022; 21(5):826–837.

Sathyamurthy A, Barik A, Dobrott CI, Matson KJE, Stoica S, Pursley R, Chesler AT, Levine AJ. Cerebellospinal neurons regulate motor performance and motor learning. Cell Rep. 2020; 31(6):107595.

Sauerbrei BA, Lubenov EV, Siapas AG. Structured variability in Purkinje cell activity during locomotion. Neuron. 2015; 87(4):840–852.

Sendhilnathan N, Semework M, Goldberg ME, Ipata AE. Neural correlates of reinforcement learning in mid-lateral cerebellum. Neuron. 2020; 106(1):188–198.e5.

Shadmehr R, Krakauer JW. A computational neuroanatomy for motor control. Experimental Brain Research. 2008; 185(3):359–381.

Snider RS, Stowell A. Receiving areas of the tactile, auditory, and visual systems in the cerebellum. Journal of Neurophysiology. 1944; 7(6):331–357.

Sobel N, Prabhakaran V, Hartley CA, Desmond JE, Zhao Z, Glover GH, Gabrieli JDe, Sullivan EV. Odorant-induced and sniff-induced activation in the cerebellum of the human. Journal of Neuroscience. 1998; 18(21):8990–9001.

Strata P. The emotional cerebellum. Cerebellum. 2015; 14(5):570–577.

Straub I, Witter L, Eshra A, Hoidis M, Byczkowicz N, Maas S, Delvendahl I, Dorgans K, Savier E, Bechmann I, Krueger M, Isope P, Hallermann S. Gradients in the mammalian cerebellar cortex enable Fourier-like transformation and improve storing capacity. eLife. 2020; 9:e51771.

Takatoh J, Prevosto V, Thompson PM, Lu J, Chung L, Harrahill A, Li S, Zhao S, He Z, Golomb D, Kleinfeld D, Wang F. The whisking oscillator circuit. Nature. 2022; 609(7927):560–568.

Tanaka H, Ishikawa T, Kakei S. Neural evidence of the cerebellum as a state predictor. Cerebellum. 2019; 18(3):349–371.

Tanaka H, Ishikawa T, Lee J, Kakei S. The cerebro-cerebellum as a locus of forward model: A review. Frontiers in Systems Neuroscience. 2020; 14.

Therrien AS, Bastian AJ. Cerebellar damage impairs internal predictions for sensory and motor function. Current Opinion in Neurobiology. 2015; 33:127–133.

Thompson RF, Steinmetz JE. The role of the cerebellum in classical conditioning of discrete behavioral responses. Neuroscience. 2009; 162(3):732–755. New Insights in Cerebellar Function.

Tseng YW, Diedrichsen J, Krakauer JW, Shadmehr R, Bastian AJ. Sensory prediction errors drive cerebellum-dependent adaptation of reaching. Journal of Neurophysiology. 2007; 98(1):54–62.

Vinueza Veloz MF, Zhou K, Bosman LWJ, Potters JW, Negrello M, Seepers RM, Strydis C, Koekkoek SKE, De Zeeuw CI. Cerebellar control of gait and interlimb coordination. Brain Structure and Function. 2015; 220(6):3513–3536.

Wagner MJ, Luo L. Neocortex-cerebellum circuits for cognitive processing. Trends in Neurosciences. 2020; 43(1):42–54.

Warren RA, Zhang Q, Hoffman JR, Li EY, Hong YK, Bruno RM, Sawtell NB. A rapid whisker-based decision underlying skilled locomotion in mice. eLife. 2021; 10:e63596.

Watson TC, Obiang P, Torres-Herraez A, Watilliaux A, Coulon P, Rochefort C, Rondi-Reig L. Anatomical and physiological foundations of cerebello-hippocampal interaction. eLife. 2019; 8:e41896.

Witter L, Rudolph S, Pressler RT, Lahlaf SI, Regehr WG. Purkinje cell collaterals enable output signals from the cerebellar cortex to feed back to Purkinje cells and interneurons. Neuron. 2016; 91(2):312–319.

Wolpert DM, Miall Rc, Kawato M. Internal models in the cerebellum. Trends in Cognitive Sciences. 1998; 2(9):338–347.

Xu-Friedman MA, Regehr WG. Ultrastructural contributions to desensitization at cerebellar mossy fiber to granule cell synapses. Journal of Neuroscience. 2003; 23(6):2182–2192.

Yang Y, Lisberger SG. Interaction of plasticity and circuit organization during the acquisition of cerebellum-dependent motor learning. eLife. 2013; 2:e01574.

